# Injectable prevascularized mature adipose tissues (iPAT) to achieve long-term survival in soft tissues regeneration

**DOI:** 10.1101/2020.12.07.415455

**Authors:** Fiona Louis, Yoshihiro Sowa, Shinji Irie, Shiro Kitano, Osam Mazda, Michiya Matsusaki

## Abstract

Soft tissue regeneration remains a challenge in reconstructive surgery. Current autologous fat implantations lead to high fat absorption ratios, while artificial implants can be associated with lymphoma occurrence. To overcome these limitations, our aim was to reproduce adipose tissue vasculature structure before implantation. Here, we developed injectable prevascularized adipose tissues (iPAT), using physiological collagen microfibers (CMF) mixed with human mature adipocytes, adipose-derived stem cells (ADSC) and human umbilical vein endothelial cells (HUVEC). Following murine subcutaneous implantation, higher cell survival (84±6% viability) and volume maintenance were shown after 3 months for the iPAT (up to twice heavier than the non-prevascularized balls). This higher survival can be explained by the greater amount of blood vessels (up to 1.6 folds increase), with balanced host anastomosis (51±1% of human/mouse lumens), also involving infiltration by the lymphatic and neural vasculature networks. These iPAT tissues allowed non-invasive soft tissue reconstruction for long-term outcomes, and the ability to cryopreserve them with maintained viability and functionality also enables a later reinjection usually required before reaching the final patient desired graft volume.

## Introduction

Adipose tissue regeneration for filling soft tissue defects has wide clinical application, affecting patients not only cosmetically (visible asymmetry or plastic surgery), but also their well-being and ability to function, such as after tumor resection, following trauma or for the treatment of high-grade burns ^1,2^. While there are many possible methods of adipose restoration, from synthetic implants to autologous fat transplantation through liposuction or deep inferior epigastric perforator (DIEP) flap methods, only natural components or the use of the patient’s own adipose tissue currently lead to acceptable results (Table 1). Artificial silicone implants in particular are associated with capsular contracture or rupture ^3–5^ and there have been recent safety concerns over their link with Breast Implant–Associated Anaplastic Large Cell Lymphoma (BIA-ALCL) occurrence (0.311 cases per 1000 person-years) ^6,7^. On the other hand, the advantages of autologous adipose tissue implantation are nevertheless tempered by its progressive resorption, by graft contracture or necrosis, possibly leading to up to 90% of volume loss and donor-site morbidity in long-term studies ^8–12^. One of the reasons is the inefficient blood supply to the fat graft, due to inadequate supportive vasculature regeneration ^13–17^. To overcome this issue therefore, the flap reconstruction has gained popularity by improving the natural aesthetic results and lowering the graft volume loss, using microsurgery to reconnect the blood vasculature *in situ*. However, the flap surgery technique requires a high skill level and long surgery time. It is also associated with morbidity, due to high risk for specific complications, such as flap necrosis from vascular anastomosis problems and blood flow insufficiency ^18^.

**Table 1:**
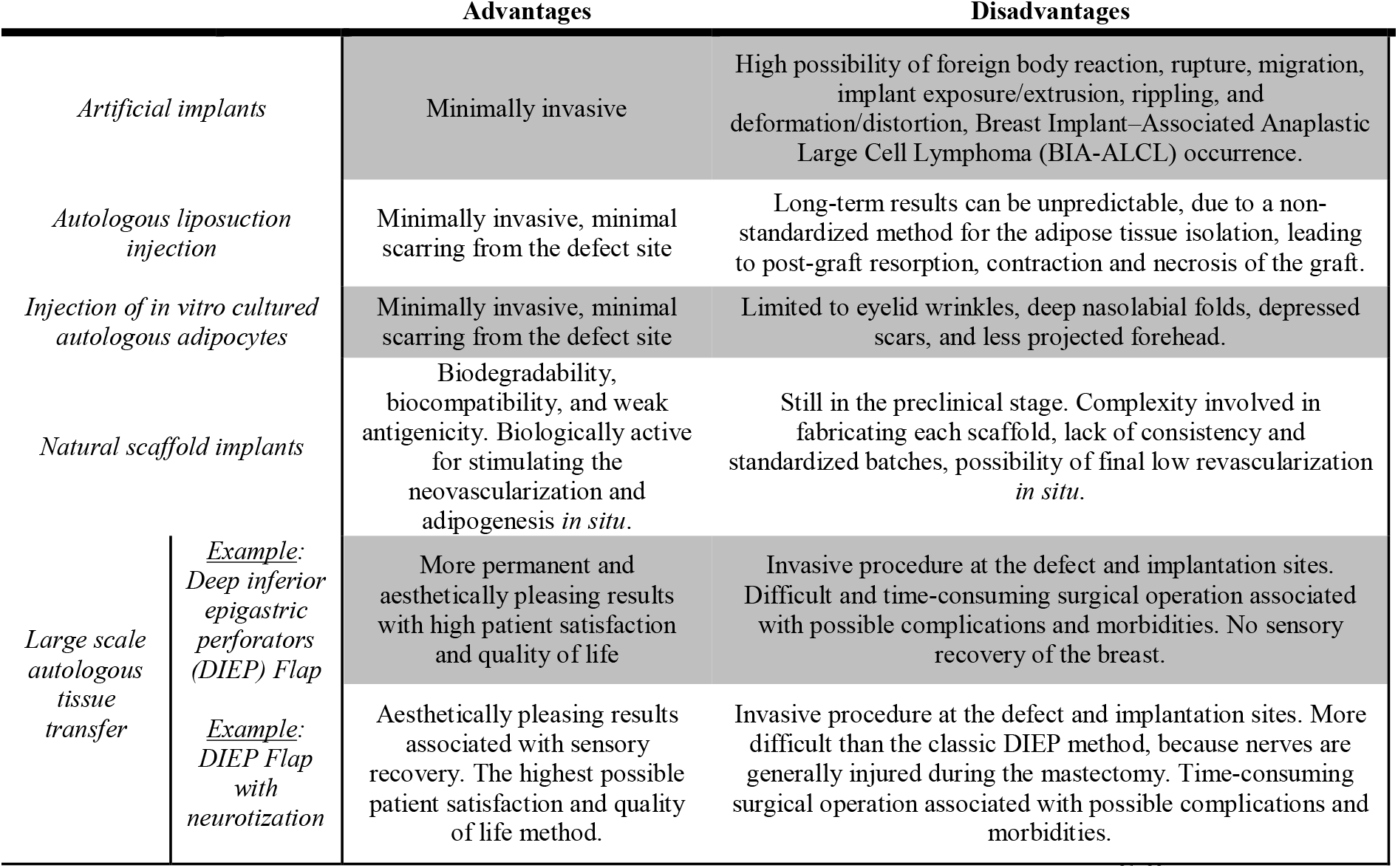
Comparison of current soft tissue regeneration surgical methods ^21–23^.

Rapid revascularization after implantation is therefore the key to achieving a successful large scale tissue graft, especially for highly vascularized adipose tissue whereby each adipocyte is normally in contact with at least one blood vessel ^19,20^. As an improvement over the current autologous liposuction injection, a versatile method with minimal invasiveness needs to be developed.

Currently, the main problems are the variations in the collection and purification of fat cells which can cause cell trauma, the dependence on centrifugation, the dimensions of aspiration and reinjection needles, and the relevance of the exposure of harvested fat to air as well as the residual infiltration solution, all of which can contribute to the low grafted adipose cell survival.^10,24,25^. In particular, mature adipocytes are greatly damaged ^8^ and stem cells or preadipocytes are not sufficiently collected ^26^, while they both help to promote the early revascularization of the autologous fat graft ^27^. One of the current strategies is thus to add adipose stem cells before transplantation (stem cell enrichment grafting like Cell-Assisted-Lipotransfer). There are some reports of positive outcomes with respect to the improved graft retention and higher capillary density found in the graft ^27,28^, but other studies have reported contrasting results ^29^.

Another point concerns the act of transplanting a large volume of adipose tissue into an enclosed space in the body which further damages the transplanted cells ^30^. Therefore, the development of an *in vitro* tissue that is resistant to mechanical stress and has a certain degree of strength would be indispensable for future improvement in fat transplantation results. This would be a more biological and practical method involving the preparation of carefully cultured, vascular-rich 3D adipose tissue containing highly active mature adipocytes and stem cells mixed in an appropriate ratio.

For effectively replicating the complexity of vascularized adipose tissue, the components used need to match the *in vivo* ones, which are mainly: mature adipocytes, adipose-derived stem cells (ADSC) and endothelial cells, in an extracellular matrix (ECM) whose main constituent is collagen type I ^31–33^. There have thus far been only two attempts to regenerate *in vitro* vascularized adipose tissues using directly mature adipocytes. Aoki et al. seeded mature adipocytes and endothelial cells from rat embedded in a collagen type I gel and found that the coculture induced the dedifferentiation of the mature adipocytes by enhancing the preadipocytes proliferation ^34^. Huber et al. performed an indirect coculture between mature adipocytes in collagen type I gel and endothelial cells ^35^. Neither of these studies showed *in vitro* blood vasculature structure formation. The induction of both adipocyte maturation and angiogenesis from stem cells remains the usual strategy, by separating the adipogenesis and the angiogenesis induction steps. Muller et al. used ADSC differentiated sequentially in endothelial cells and then adipocytes using spheroids culture embedded in Matrigel^® 36^. Hammel et al. took the opposite route by first differentiating human mesenchymal stem cells (hMSC) into adipocytes before adding human umbilical cord vein endothelial cells (HUVEC) in a collagen type I gel ^37^. Kang et al. also cultured ADSC and then HUVEC on a porous silk protein scaffold ^38^. Qi et al. used several synthetic hydrogels (methacrylated gelatin, methacrylated hyaluronic acid and 4arm poly(ethylene glycol) acrylate (PEG-4A)) for the culture of ADSC and then HUVEC ^39^, while Paek et al. used ADSC and then human adipose microvascular endothelial cells (hAMECs) in a hydrogel scaffold composed of type I collagen and fibrin ^40^. Finding the appropriate culture medium to enable both the differentiation of adipocytes up to their mature differentiated state and the endothelial cells vasculature formation in coculture could still not be achieved in any of these models however.

To counteract these problems, we recently developed collagen microfiber (CMF) tissues which improved the maintenance of mature adipocyte viability and functions ^41^. Unlike classic collagen gel using dilution, these dispersed collagen microfiber scaffolds allowed the attainment of a final physiological concentration closer to the one found in adipose tissue ^42^, while protecting the fragile mature adipocytes from shear stress. The endothelial cells were also found to be able to attach to these collagen microfibers, helping the vasculature construction through integrin-induced adhesion ^43,44^.

The purpose of this study was thus to construct injectable prevascularized adipose tissues (iPAT) as *in vivo*-like adipose ball-shaped tissues containing: mature adipocytes, ADSC and HUVEC, mixed in a fibrin gel matrix (Figure 1). Then we implanted them subcutaneously in mice and the grafted iPAT outcomes showed improved survival compared to the implantation of engineered adipose ball tissues without prevascularization or the classic liposuctioned adipose tissue. The key advantage of this method relied on the higher cell engraftment, survival and tissue volume maintenance, probably due to a stronger host-vasculature connection also involving the lymphatic and neural vasculature networks. Another benefit of this technology was also the ability to store the iPAT balls made from only one liposuction operation, allowing the later non-invasive reinjection of iPAT if needed for final patient’s graft volume correction.

**Figure 1:**
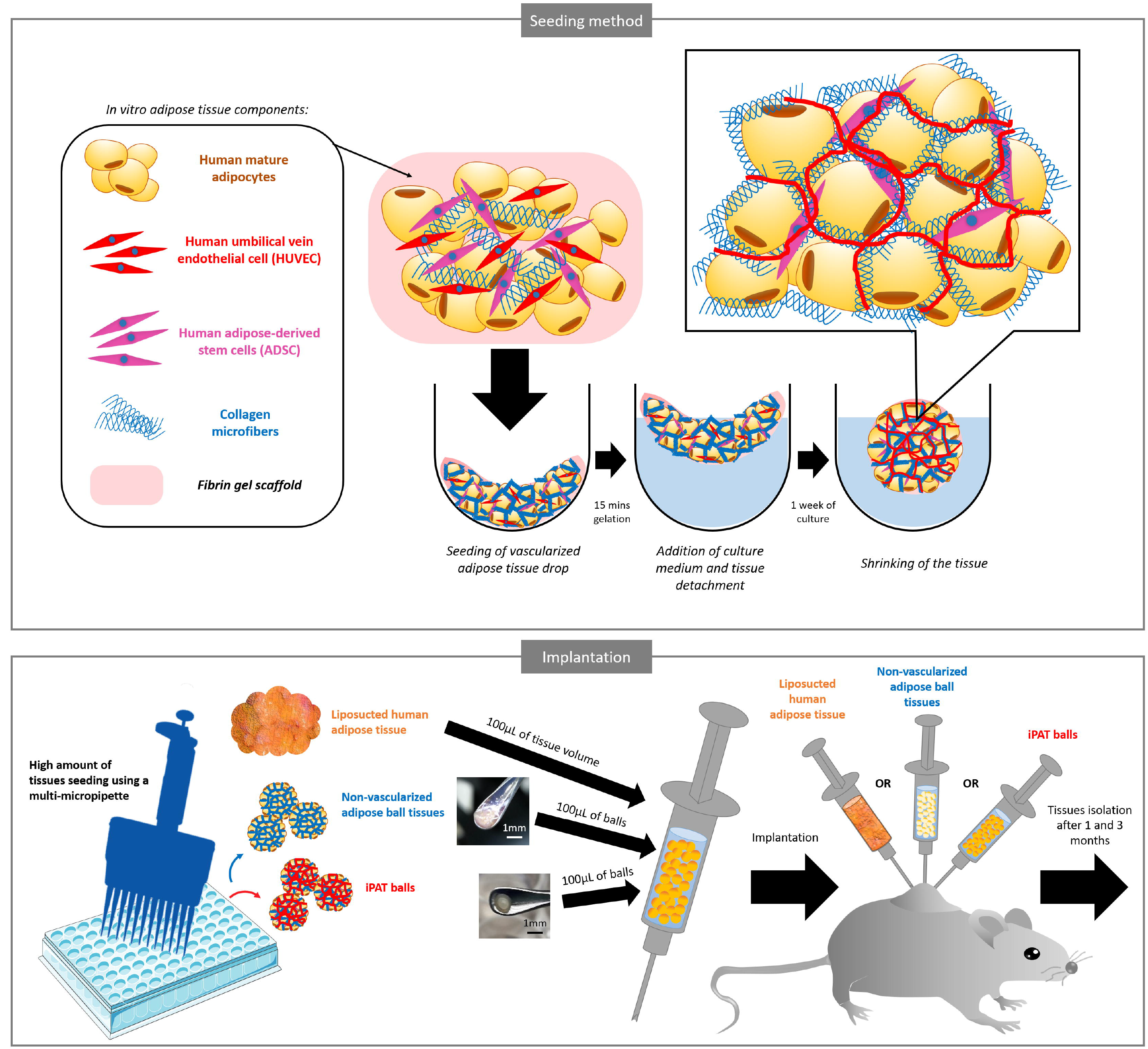
Injectable prevascularized adipose tissues (iPAT) construction and implantation process. Injectable prevascularized adipose tissue (iPAT) balls were composed of human mature adipocytes, human umbilical vein endothelial cell (HUVEC) and human adipose-derived stem cells (ADSC), mixed with collagen microfibers type I and embedded in a fibrin gel. A drop of this mixture was seeded on a non-adherent plate and after gelation, culture medium was added, leading to drop detachment and its shrinking, to make a ball shape after 7 days of culture. Non-vascularized ball tissues were also produced using the same method but without HUVEC and ADSC. Using a multi-micropipette, a large number of ball tissues were seeded at the same time in order to get 100µL of final volume to inject in the mouse. A control sample consisted of classic liposuctioned human fat tissue. Each different condition was injected on one side of the mice’s back for tissue analysis.

## Results

### *In vitro* prevascularized adipose tissue model reconstruction

While mature adipocytes and ADSC can be obtained easily and in large amounts by isolation from patients’ adipose tissues, mature adipocytes still remain difficult to maintain in culture ^45^. The first step to be able to reconstruct *in vitro* prevascularized adipose tissue was thus to check their viability when using CMF (Figure 2a). Live/dead images showed that mature adipocytes mixed with CMF can keep their unilocular lipid vesicle phenotype, with only a few spindle-shapes dedifferentiated cells, for up to three weeks of *in vitro* culture. This high viability was found to differ non-significantly at 94±7%, 90±3% and 89±3% for days 7, 14 and 21 of culture respectively. The second step was to add other adipose cell types in coculture until finding the right mixture allowing the vasculature formation from HUVEC, monitored by the CD31 endothelial marker immunostaining. Generally, mesenchymal cells like fibroblasts are required to secrete the growth factors cocktails necessary for vasculature network initiation ^46^. As ADSC derive from the same mesenchymal progenitor, they should also induce angiogenesis ^47,48^. In this study, while mature adipocytes cocultured with HUVEC showed a limited vasculature (Supplementary Figure 1a, white arrows), it did not appear when normal human dermis fibroblasts (NHDF) were also added. Only ADSC and HUVEC coculture allowed a dense vasculature formation, which was even more strongly induced when mature adipocytes were also present, surrounded by capillaries in the same way as *in vivo*.

**Figure 2:**
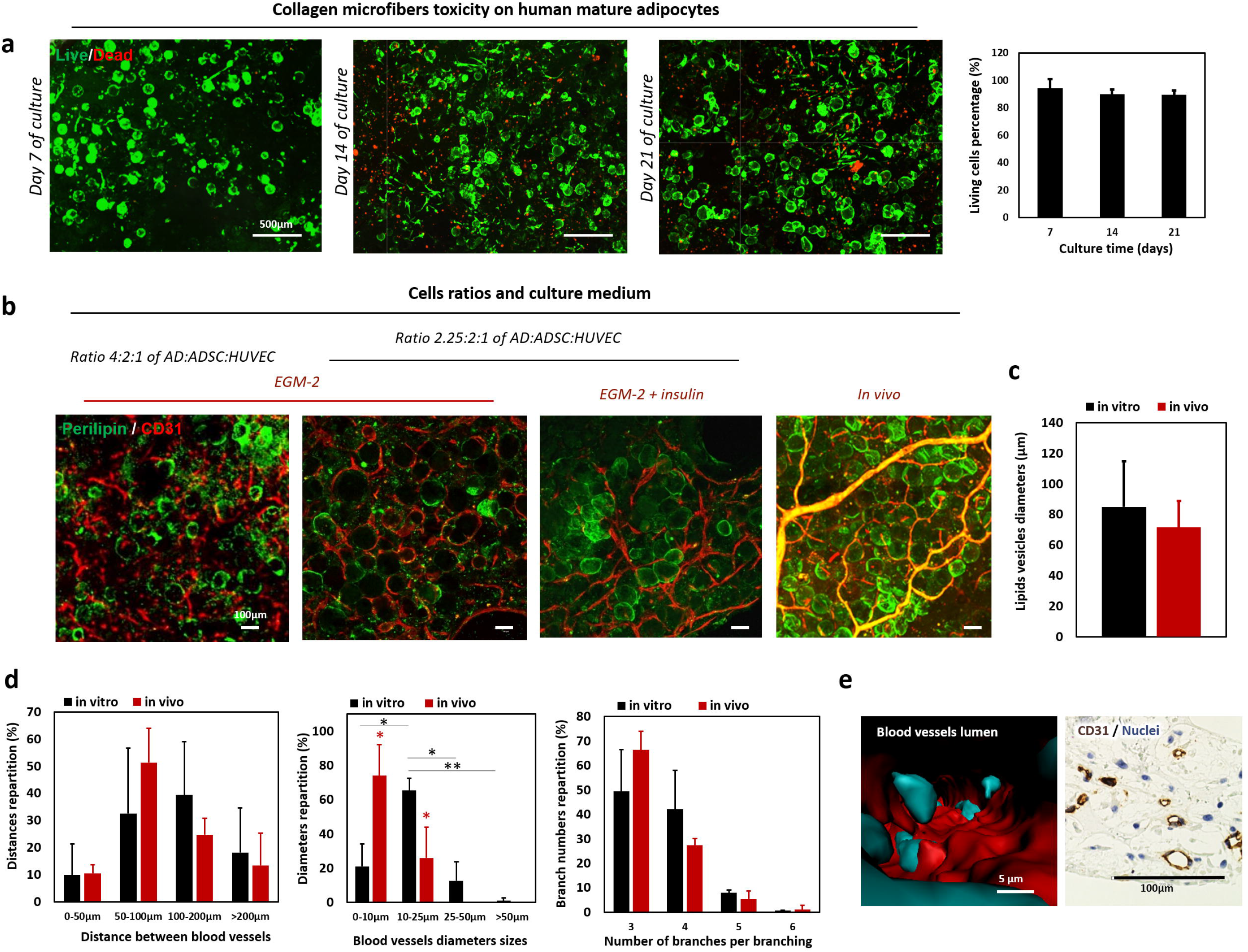
Preliminary developments for a suitable prevascularized mature adipose tissue induction *in vitro*. (a) Representative Live-Dead projection images of mature adipocytes cultured in collagen microfibers (CMF) up to 21 days. Live/Dead quantitation, results are means□±□s.d. on n=3 independent samples, 6 pictures measured. (b) Representative immunostaining images for perilipin of lipid vesicles and CD31 of blood vessel vasculature depending on the cell ratios and the culture medium components, compared to an *in vivo* image (Ratio 4:2:1 is 8×10^6^ mature adipocytes/mL + 4×10^6^ ADSC/mL + 2×10^6^ HUVEC/mL; ratio 2.25:2:1 is 4.5×10^6^ mature adipocytes/mL + 4×10^6^ ADSC/mL + 2×10^6^ HUVEC/mL). (c) Lipid vesicle diameter measurements of mature adipocytes in the EGM-2+insulin 10µg/mL (EGM-2 being the endothelial medium) *in vitro* condition compared to the native (*in vivo*) tissues. The results are means□±□s.d. of experiments performed on three independent samples, n=35-62 adipocytes measured. (d) Measurements of their diameter repartitions (n=56-168 diameter measures per experiment, three independent experiments, \**p*<0.05 and \*\**p*<0.01), the distance between the blood vessels (n=189-256 distances measured per experiment, three independent experiments), and the number of branches per branching (n=48-149 branching measured per experiment, three independent experiments) in the *in vitro* tissues compared to the *in vivo* condition (3 different donors). The results are means□±□s.d. (e) Representative Imaris software 3D reconstruction image from CD31/Hoechst immunostained samples showing inside a lumen in a blood vessel of an *in vitro* tissue, and representative histology paraffin section of CD31 immunostaining showing the lumens inside the *in vitro* tissues.

Next, using these three types of cells, the effect of the scaffold matrix components for the vasculature formation optimization was assessed (Supplementary Figure 1b). CMF alone was already found to be associated with the initiation of several blood vessel structures but the observed high shrinkage of the tissue prevented the construction of a homogeneous organization of capillaries. Only the condition at 1.2wt% of final CMF concentration, mixed with fibrin gel, allowed a dense vasculature network formation throughout the tissue. The use of only fibrin gel or three times more concentrated CMF (3.6%wt) conditions both inhibited the structural formation of blood vessels.

Finally, the optimization of the cell ratios, as well as the culture medium conditions were performed (Figure 2b). By seeding twice as many mature adipocytes as ADSC and four times as many as HUVEC, the vasculature formation was found altered with few continuous vessels surrounding the mature adipocytes, stained by the lipid vesicle perilipin marker, compared to a reduced mature adipocyte concentration which showed a dense CD31 network. As until this point only an endothelial medium (EGM-2) had been used to induce the vascularization during *in vitro* culture, the culture medium was also optimized to obtain a better maintenance of the mature adipocytes which sometimes displayed an altered perilipin staining. By adding insulin at 10µg/mL in the EGM-2 medium, all of the mature adipocytes maintained their unilocular lipid vesicle throughout the culture, with a surrounding dense blood vessel network, to the same extent as in *in vivo* human adipose tissue. Effective maintenance of the unilocular phenotype was confirmed by the measurement of 85±30µm for their lipid vesicle average diameter, differing non-significantly from *in vivo* native mature adipocytes (71±17µm average diameter, Figure 2c), while the viability of the whole tissue was found to be high (85%±8, Supplementary Figure 2), even if it was difficult to distinguish mature adipocytes from the other cell types. A comparison of the distances measured between each blood vessel in the tissues, *in vivo* and *in vitro*, yielded no significant differences (Figure 2d), the averages being 135±55µm *in vitro* and 120±84µm *in vivo*. Most of the distances (72% for *in vitro* and 76% for *in vivo*) were found to be between 50 and 200µm which is exactly in the range of diameter of one human mature adipocyte (as seen in Figure 2c), reinforcing the fact that as *in vivo*, all mature adipocytes in this model seemed connected to at least one blood vessel. On the other hand, the vessel diameters repartition revealed larger CD31+ vasculature *in vitro* compared to *in vivo*, with 65% of *in vitro* vessel diameters being between 10 and 25µm (small venules) while 74% of the *in vivo* vessel diameters were between 0 and 10µm (capillaries), for an average diameter of 16±4µm and 8±4µm for *in vitro* and *in vivo* vessels respectively. The branch numbers repartition in the vasculature structures were also compared and did not show any significant differences between *in vitro* and *in vivo*. Finally, to validate the ability of the observed *in vitro* blood vessels to allow fluid diffusion, lumen structures were observed inside the samples, by CD31 3D reconstruction analysis images and CD31 immunohistochemistry on paraffin slides, which should allow the nutrient diffusion through the tissues (Figure 2e).

### Injectable prevascularized adipose tissues (iPAT) reconstruction

After validating the right condition for achieving dense vascularization around the mature adipocytes in *in vitro* culture, the same mixture was proceeded to design adipose ball tissues that can be injected and merged to form a larger tissue *in situ*: the injectable prevascularized adipose tissues (iPAT). The first step was to compare different seeding volumes to find a final ball tissue diameter suitable for needle aspiration and injection, thus avoiding invasive implantation surgeries for the patients (Figure 3a). Three seeded volumes were compared: 5, 10 and 15µL. After 7 days of culture, the diameters of the vascularized ball tissues were 1,250±118µm, 1,829±159µm and 2,165±296µm, depending on the seeded volume, for 5, 10 and 15µL respectively. The 5µL volume was thus chosen as being suitable for the iPAT balls, in order to be able to aspirate the ball tissues using a 15G needle on a syringe (needle inner diameter of 1,370µm). The obtained iPAT balls showed the same dense blood vessel vasculature after 7 days of culture, surrounding the mature adipocytes (Figure 3b), as seen in the transwell tissues structure shown in Figure 2. These blood vessels were found around but also inside the balls as confirmed by the CD31 staining on their paraffin sections, with some lumen structures observed, as well as blood vessels surrounding the mature adipocytes (Figure 3c).

**Figure 3:**
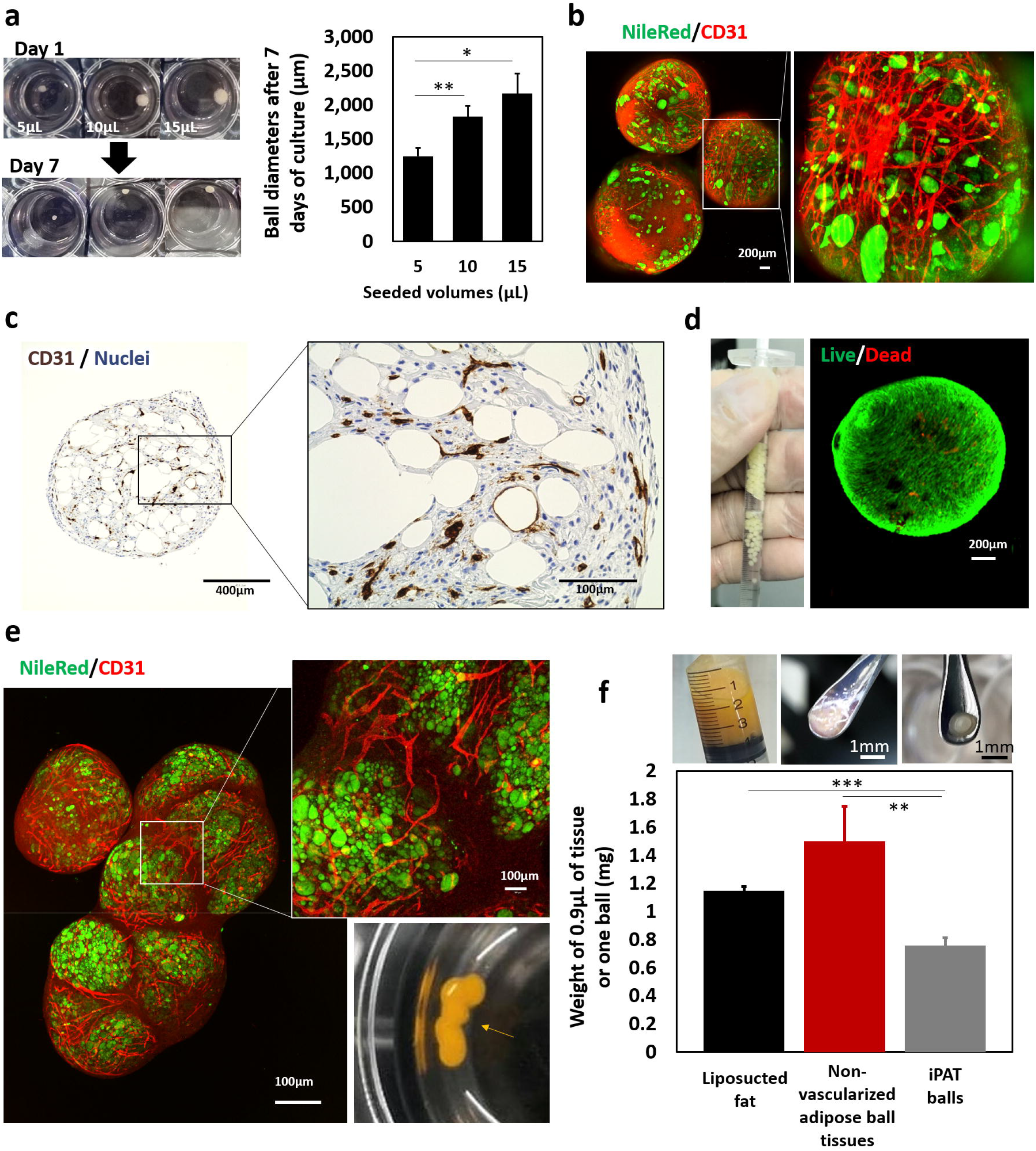
Adipose ball tissues construction, injection validation and ball merging assessment before implantation. (a) iPAT ball tissue images of 5, 10 and 15 µL volumes shrinkage during the 7 days of culture and measurement of the tissue diameters after 7 days. The results are means□±□s.d. of n=12 balls per condition, \**p*<0.05 and \*\**p*<0.01. (b) Representative images of NileRed staining for lipid vesicles and CD31 immunostaining for blood vessel vasculature of the iPAT tissues. (c) Representative histology paraffin section of CD31 immunostaining showing the lumens inside the iPAT tissues. (d) Validation of injection feasibility using a 15G needle on a 1mL syringe. Representative image of the Live/Dead staining of the ball tissues after their several aspiration and release steps using the syringe. (e) Representative images of NileRed staining for lipid vesicles and CD31 immunostaining for blood vessel vasculature after an additional week of culturing several iPAT tissues in the same culture well to obtain a higher size tissue by the merging of balls. Vasculature connection can be observed between the merged balls. (f) Liposuctioned human adipose tissue, non-vascularized adipose ball tissue and iPAT ball tissue pictures. Measured weights of the three types of tissues corresponding to one ball or 0.9µL equivalent volume (measures realized three times on 3x 10 different balls per condition). Graph results show means□±□s.d., \*\**p*<0.01 and \*\*\**p*<0.001.

The ability to aspirate the iPAT balls in the syringe was then assessed (15G needle on a 1mL syringe). After 7 days of culture, the balls were aspirated and released three times and their viability was assessed 24 hours after, showing only a few dead cells (Figure 3d). Next, the confirmation of the ability of the iPAT balls to merge and create a customized higher final tissue size after implantation was monitored. Several balls were cultured together in the same culture well for an additional 7 days (totally 14 days of culture). The result showed five aggregated balls (Figure 3e). In the merged balls, the vasculature network was still observed after the additional 7 days and the blood vessels were even found connected between the different balls, which should ensure adequate nutrient and oxygen diffusion even in the large final tissue size. Another check was to see if an adipose ball tissue embedded in fibrin gel, as it will be implanted *in vivo*, can expand its blood vessels to ensure proper nutrient and oxygen supply before the fibrin gel degrades. After an additional 7 days in culture, CD31 immunostaining of the embedded balls showed blood vessels reaching the outside of the ball as shown in Supplementary Figure 3.

The injection process through the skin itself was then checked by injecting non-vascularized adipose balls and iPAT balls made from the cells of the same patient as the skin parts (Supplementary Figure 4). The 7-days ball tissues were injected into a fibrin gel below the skins and kept an additional week in culture as well. By observing from the bottom of the transwells containing the skin, the iPAT blood vessels were found extended and in connection with the skin blood vessels. This connection was less evident with the non-vascularized adipose ball tissues. Finally, Figure 3f shows the three types of tissues to be implanted *in vivo*: the “liposuctioned” human adipose tissue from which 100µL was injected below the mice back skin, the non-vascularized adipose balls and the iPAT balls, of which 111 were subcutaneously injected, to get a final implanted volume of 100µL. iPAT showed a smaller diameter and a spherical shape, while non-vascularized balls displayed a flat shape. The weight of 0.9µL of the three types of tissues (corresponding to one iPAT’s volume) was also measured and compared. The non-vascularized balls displayed the highest weight with an average of 1.5±0.3mg per ball, possibly due to the higher water content in each ball. The liposuctioned fat and the iPAT showed lighter average weights of 1.1±0.03mg and 0.8±0.06mg respectively. These weights will be useful for the comparison of the survival weights after *in vivo* implantation.

### *In vivo* mice implantation survival and tissue weight retention results after 1 and 3 months

The mice were implanted with two tissues, each tissue being of a different condition (iPAT, non-vascularized adipose ball tissues or the control: liposuctioned adipose tissue), in order to get the three possible combinations, and the tissues were isolated after 1 and 3 months. Figure 4a shows the isolation of the tissues from the mice, some tissues being visible below the skin. The individual balls forming the merged iPAT can still be observed in the final tissue after isolation (Figure 4a right). Some oily/transparent parts can sometimes be seen on some tissues (Figure 4b + all the tissues isolated from the three independent experiments in Supplementary Figure 5), especially in the liposuctioned and non-vascularized samples. These areas might reveal necrotic parts, as can be observed in Figure 4c, where empty areas are observed on the H/E staining histology sections of the tissues for these two conditions to a much greater extent than in the iPAT ones (see the asterisks). The H/E staining also allowed us to observe some red blood cells inside the lumens, for the three conditions, testifying to the blood perfusion of the tissues from the host vasculature connection (Figure 4c, right).

**Figure 4:**
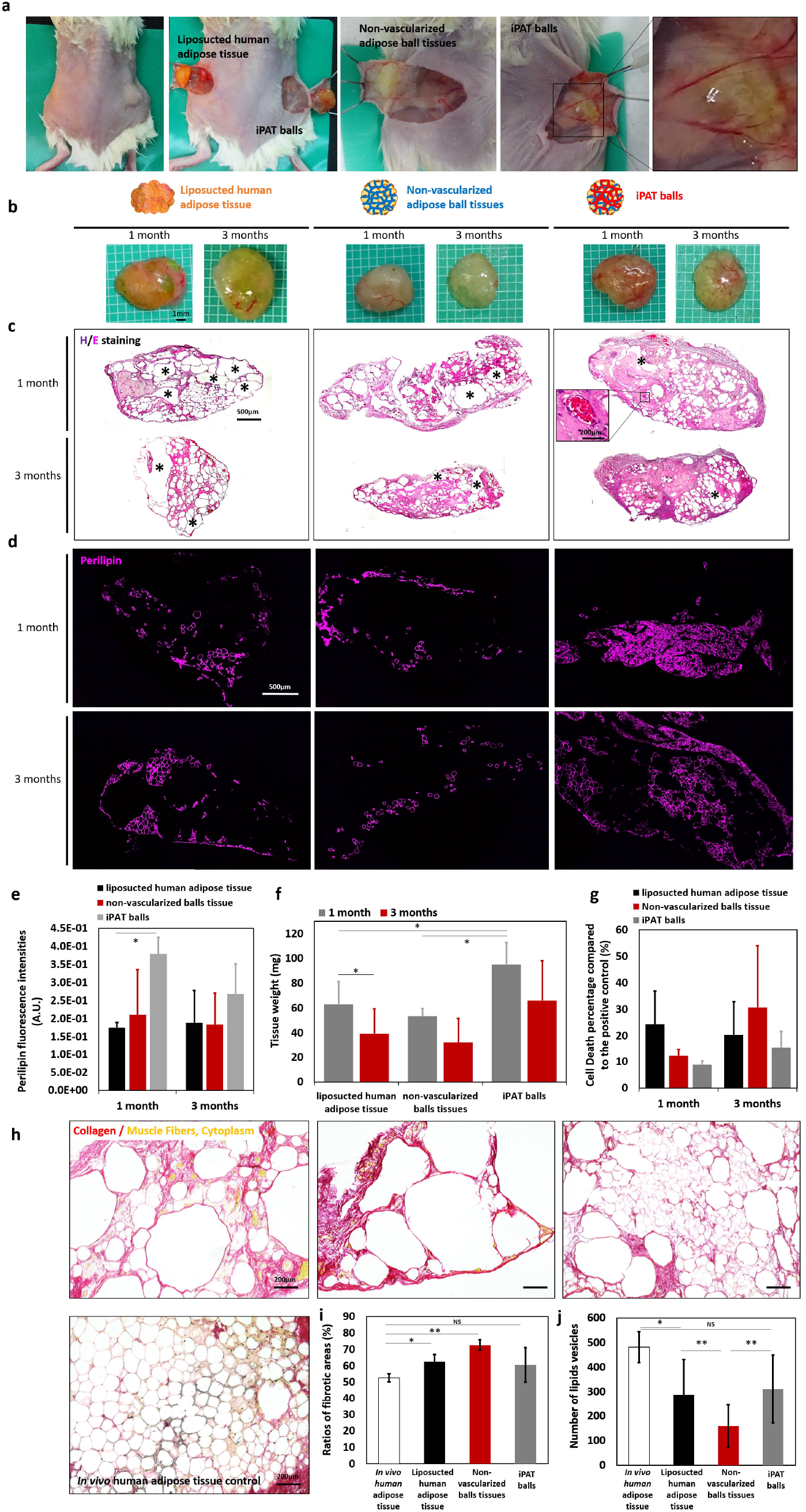
*In vivo* mice implantation survival and tissue volume maintenance results after 1 and 3 months. (a) Representative image of an implanted mouse after 1 month. Both sides tissue isolation after mouse euthanasia. Individual adipose ball tissues can still be observed from the merged final tissue. (b) Representative isolated tissue pictures for the three conditions, after 1 and 3 months implantation in mice. (c) Representative histology paraffin section of Hematoxylin/Eosin (H/E) staining on the ball tissues for the three conditions after 1 and 3 months of implantation. Asterisks represent the necrotic areas found inside the tissues. (d) Representative images of perilipin immunostaining of lipid vesicles in the tissues for the three conditions after 1 and 3 months of implantation. (e) Quantification of perilipin fluorescence intensities in the centre of the grafts, in each condition. (f) Tissue weight measurements in each condition and depending on the implantation time. (g) Cell death assessment measurement quantification for the three conditions. (h) Representative images of the elastica van Gieson (EVG) staining of the tissues. Collagen (red) and muscle fiber or cell cytoplasm (yellow) allow the estimation of fibrotic percentage. (i) Quantification of fibrotic area ratios in each conditions and compared to *in vivo* human adipose tissue. (j) Quantification of lipid vesicle number density in the tissues, in each condition. All graph results show means□±□s.d. of the three independent experiments performed. Measurements were performed on n=3-5 sections images per sample. \**p*<0.05 and \*\**p*<0.01.

Adipocyte-specific perilipin immunostaining was then performed on the paraffin sections (Figure 4d). Compared to traditional ‘‘all-or-nothing’’ markers of viability (phycoerythrin or trypan blue), a declining perilipin expression can indicate here a declining viability, sometimes found in implanted grafts and initiated in their centre, where both adipocytes and ADSC prematurely die (necrotic zone) ^10^. As seen in the pictures, mature adipocytes were still found in the implanted tissues, even after 3 months *in vivo*, and perilipin staining was found to be significantly higher in the centre of the graft for the iPAT compared to the two others especially after 1 month of implantation (2.2 and 1.8 times more than the liposuctioned fat and non-vascularized ball tissues respectively, Figure 4e).

To further compare the different tissues, their weights were measured (Figure 4f) and appeared significantly reduced in the liposuctioned (1.5 times lighter) and non-vascularized ball tissues (1.8 times) conditions, when compared to the iPAT balls after 1 month, and non-significantly reduced after 3 months, the iPAT balls being still 1.7 and 2 times heavier than the liposuctioned fat and non-vascularized ball tissues respectively.

To better understand this difference in the tissue volume retention, their cell death resulting from both necrosis and apoptosis was assessed (Supplementary Figure 6 and quantitation in Figure 4g). Surprisingly, a high cell death was seen after 1 month first on the liposuction injected tissues (24±12%), while after 3 months it was found more in the non-vascularized ball tissues (31±23% of cell death). The iPAT were associated with the highest cell survival after both 1 and 3 months (91±1% and 84±6% respectively), which could explain their higher weight retention as seen in Figure 4f. Another crucial assessment for the clinical application, linked to the cell survival and the occurrence of necrosis, was to monitor the ratio of the fibrotic areas after the 3 months of mice implantations, by using EVG staining to visualize elastic and collagen fibers (Figure 4h-i). When compared to *in vivo* human adipose tissue, liposuctioned fat injection and non-vascularized ball tissues showed significantly higher fibrotic area ratios (respectively +10 and +20%), which was not the case for the iPAT. As for elastic fibers, it appeared that the three injected conditions no longer contained any, compared to the *in vivo* adipose tissue (black staining). The elastic fiber content strongly varies depending on the location and the patient condition (lean or obese), it could have come from a case with few elastic fibers or it may be due to the fact that the blood vessels in the *in vitro* adipose tissues are still immature compared to *in vivo* where blood vessels are surrounded by elastic fibers. As overly large lipid vesicles can also indicate mature adipocyte necrosis, it was important to finally check the lipid vesicles’ size and density by calculating the lipid vesicle number per sectioned tissue area (Figure 4j), a low lipid vesicle density suggesting fat necrosis or fibrosis. Again, non-vascularized adipose ball tissues were associated with a significantly lower density compared to the liposuctioned fat and iPAT (−44% and −49% respectively), even though both of them were also found to be lower than the *in vivo* fat lipid vesicles density (−40% and −35% respectively).

### Blood vessel monitoring after implantation

To more deeply understand the iPAT’s higher survival, the vascularization in the tissues was monitored after 1 and 3 months of *in vivo* implantation. First, the von Willebrand factor (VWF) endothelial marker area was assessed and quantified after the 3 months of implantation. It was found to be significantly lower especially in the non-vascularized adipose ball tissues (Figure 5a), while liposuctioned fat and iPAT showed no differences compared to *in vivo* adipose tissue. CD31 endothelial marker was also observed and quantified depending of the conditions and the implantation time (Figure 5b + Supplementary Figure 7). While lumens were displayed in the three conditions, the stained areas were significantly larger in the iPAT compared to the two other conditions both after 1 and 3 months of implantation (3.4 and 2.3 fold increases at 1 month, 1.6 and 1.4 fold increases at 3 months compared to liposuctioned fat and non-vascularized adipose ball tissues respectively).

**Figure 5:**
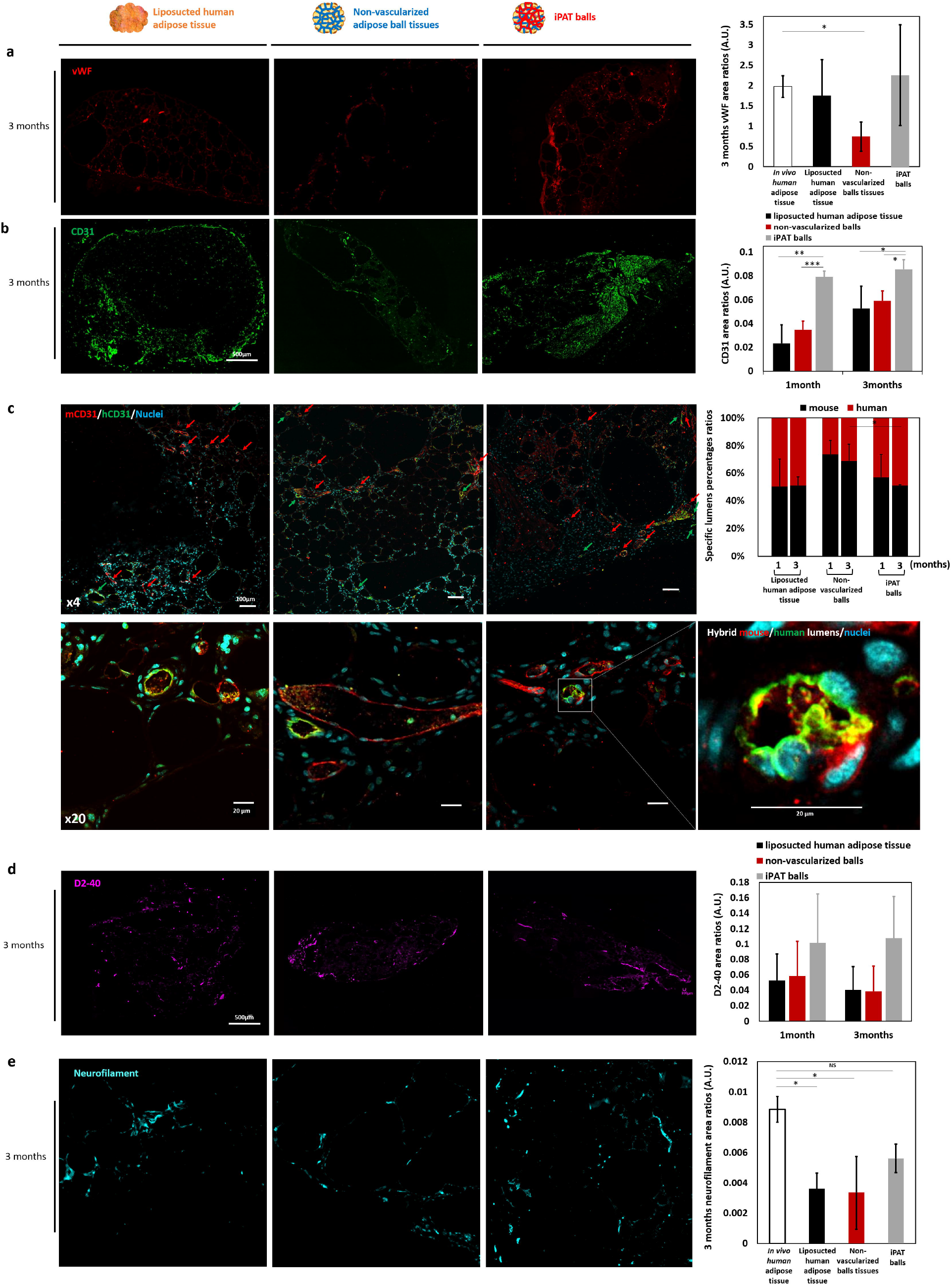
Blood and lymphatic vasculature as well as neural network assessments of the implanted tissues. (a) Representative images of vWF endothelial marker immunostaining in the tissues for the three conditions after 3 months of implantation. Stained area ratios quantification in each condition and the implantation duration, compared to the *in vivo* human adipose tissue. (b) Representative images of combined mouse and human CD31 immunostainings of blood vessels in the tissues for the three conditions after 3 months of implantation. Stained area ratios quantification regarding the conditions and the implantation duration. (c) Representative images of specific mouse or human CD31 immunostainings in the tissues for the three conditions after 1 and 3 months of implantation. Green arrows represent the human lumens observed and red arrows the mouse ones. Quantification of the specific lumen ratios regarding the species and according to the conditions. Enlargement of a representative hybrid lumen from the vascularized adipose ball tissues showing both human and mouse endothelial cells on the same lumen. All graph results show means□±□s.d. of the three independent experiments performed. Measures were performed on n=3-5 sections images per samples. \**p*<0.05 and \*\**p*<0.01. (d) Representative images of D2-40 immunostainings of mouse lymphatic vessels in the tissues for the three conditions after 3 months of implantation. Stained area ratios quantification regarding the conditions and the implantation duration. (e) Representative images of neurofilament immunostainings in the tissues for the three conditions after 3 months of implantation. Stained area ratios quantification regarding the conditions and the implantation duration, compared to *in vivo* human adipose tissue. All graph results show means□±□s.d. of the three independent experiments performed. Measures were performed on n=3-5 sections images per samples. \**p*<0.05 and \*\**p*<0.01.

The lumens were then further inspected. After the mice implantation of the reconstructed tissues using human cells, both types of lumen could be observed, coming from mouse (host) or human (graft) origins. Specific CD31 immunostainings were thus performed (Figure 5c), showing both species types (human lumens by green arrows and mouse lumens by red arrows) in the three conditions, with sometimes even hybrid lumens formed by both mouse and human endothelial cells observed, demonstrating the anastomosis of the engineered vessels with the host’s vasculature. The number of mouse or human lumens in the same area was then counted, and the results of the average of the three experiments showed a significantly higher ratio of mouse lumens (74±10% and 69±12% at 1 and 3 months) in the non-vascularized adipose ball tissues especially at 3 months, while balanced ratios appeared in the two other conditions at 1 and 3 months (50±20% and 50±6% for liposuctioned tissues, 57±17% and 51±1% for iPAT). For the non-vascularized adipose tissues samples, only human mature adipocytes were injected in the mouse. However, during the mature adipocytes isolation process, a very small amount of other cell types (endothelial cells or ADSC) from the adipose tissue can stay attached to the mature adipocytes (around 0.2-0.4% generally ^34,49^) and can then expand in the *in vitro* balls culture as well as after implantation. Another explanation is that these human endothelial cells came from the dedifferentiation of the mature adipocytes which can then transdifferentiate in endothelial cells ^50^. For the liposuctioned tissues and the iPAT, equal ratios were observed but different cell death results (Figure 4g) with a higher cell death found after 1 month in the liposuctioned fat condition. This could be due to the fact that the connection between the human blood vessels of the implant and the host blood vessels was not yet effective after 1 month, leading to the necrotic areas found in Figure 4c. In comparison, the overly high ratio of mouse blood vessels observed in the non-vascularized adipose ball tissues could provide an explanation for the high cell death found after 3 months. This could be linked to the difficulty of mouse blood vessels to penetrate deeply into the implanted tissue, staying more on the surface and impairing the nutrient and oxygen diffusion through the whole graft. In fact, the CD31 immunostaining images of the full tissue effectively showed a low number of lumens in the middle of these tissues after 3 months (Figure 5b).

### Lymphatic vasculature and neural network assessments of the implanted tissues

The higher cell survival and tissue volume retention could also be explained by the lymphatic vasculature network, in addition to the blood vessel network. D2-40 immunostaining, a specific marker of lymphatic endothelial cells, was thus performed on sectioned tissues and the stained areas were quantified (Figure 5d + Supplementary Figure 8). No significant differences were seen after 1 month and 3 months of implantation with lymphatic vessels found on the tissue edges, but iPAT tissues were found to have a higher tendency of lymphatic vessels tissue invasion at both times.

This last result, combined with the balanced ratio of mouse and human lumens could be the key to higher adipose tissue survival through a successful anastomosis with the host’s vasculature after *in vivo* implantation.

A final important check was on the neurofilaments, intermediate filaments found in the cytoplasm of neurons, indicating the neurons’ invasion in the implanted tissues (Figure 5e + Supplementary Figure 9a-b). Knowing that fat storage and metabolism are also regulated by neural factors to ensure a proper adipose tissue homeostasis and thus its survival ^51^, the better implanted tissue survival found in the iPAT can also be explained by the higher neural invasion found, compared to the liposuctioned and non-vascularized adipose ball tissues conditions. This neural network can come from the host or already be included in the ball tissues before implantation. To confirm this last point, the 7-day culture ball tissues were also immunostained for the neurofilaments before their implantation (Supplementary Figure 9b). Only the iPAT presented some neurofilaments, following some capillaries of the balls’ blood vessels vasculature. The alignment of peripheral nerves with blood vessels, forming the intricate branching networks that connect every organ, actually relies on the vascular tree to supply the necessary oxygen and nutrients ^52^. The vascular endothelial cells can guide the regeneration of peripheral nerve axons by their production of vascular endothelial growth factor (VEGF) which also stimulates the outgrowth of Schwann cells and enhances axons ^53^. These neural cells observed in the iPAT, helping the implant to connect with the host neural network, can be originated from the adipose cells isolation which contains Schwann cells ^54^, or from the ADSC, their neural transdifferentiation having already been observed in *in vitro* study ^55^.

### Feasibility of cryopreservation storage of the injectable ball tissues

One of the main obstacles of the current human liposuction and reinjection process for breast reconstruction is that it generally needs overcorrection and reinjection procedures to achieve long-term favourable results due to its high-volume loss, also depending on the morphology of the tissue defects. This means that repeated liposuction operations are required, implying higher costs, patient morbidity, and discomfort. As it cannot be determined in advance what the final volume as a long-term outcome of one implantation will be, to maximize the extracted fat tissue from liposuction surgery, it is important to be able to transplant iPATs by multiple operations until the desired volume is attained. To address this concern, we also checked the feasibility of cryopreserving the ball tissues of several prior *in vitro* cultures from the same patient cells, to be able to perform reinjection later. Generally, the mature adipocytes’ fragility makes it difficult for them to survive after freezing/thawing steps. A cryopreservation assay was thus performed on the adipose ball tissues and their viability was checked after 7 and 30 days of freezing storage (Figure 6a-b). The iPAT balls showed a highly preserved viability after 7 and 30 days, which was not significantly different to day 0 before the cryopreservation, while the non-vascularized balls had significantly lower viability after 7 days at −160°C (73±13% and 59±4% of living cells for the LaboBanker and the Trehalose cryopreservative media conditions respectively) and 30 days (72±6% and 59±2%). It seems that the added cell types (ADSC and HUVEC), in addition to the protective shrunken ball shape, improved the survival of the tissues following their storage. Another check, in addition to the viability assessment, was to monitor the maintained functionality of the tissue, especially for the mature adipocytes, known for their fragility following freezing storage. First, the iPAT balls’ ability to merge was confirmed when cultured together during 7 days even after 30 days of freezing storage, showing the maintenance of the blood vasculature connection between the ball tissues (Figure 6c). Then, one function of the mature adipocytes, their fatty acids uptake, was monitored using the fluorescently tagged fatty dodecanoic acid (C12H24O2) analog (BODIPY™ 500/510 C1, C12), after insulin induction. This non-invasive assay assessed the fatty acid accumulation in the unilocular intracellular lipid vesicles after 60 minutes (Figure 6d). These results confirmed the feasibility of storing the iPAT balls for patients, constructed from one liposuction operation, in order to provide a stock which can be used to readjust the injected graft volume by subsequent injections depending of the volume loss outcomes.

**Figure 6:**
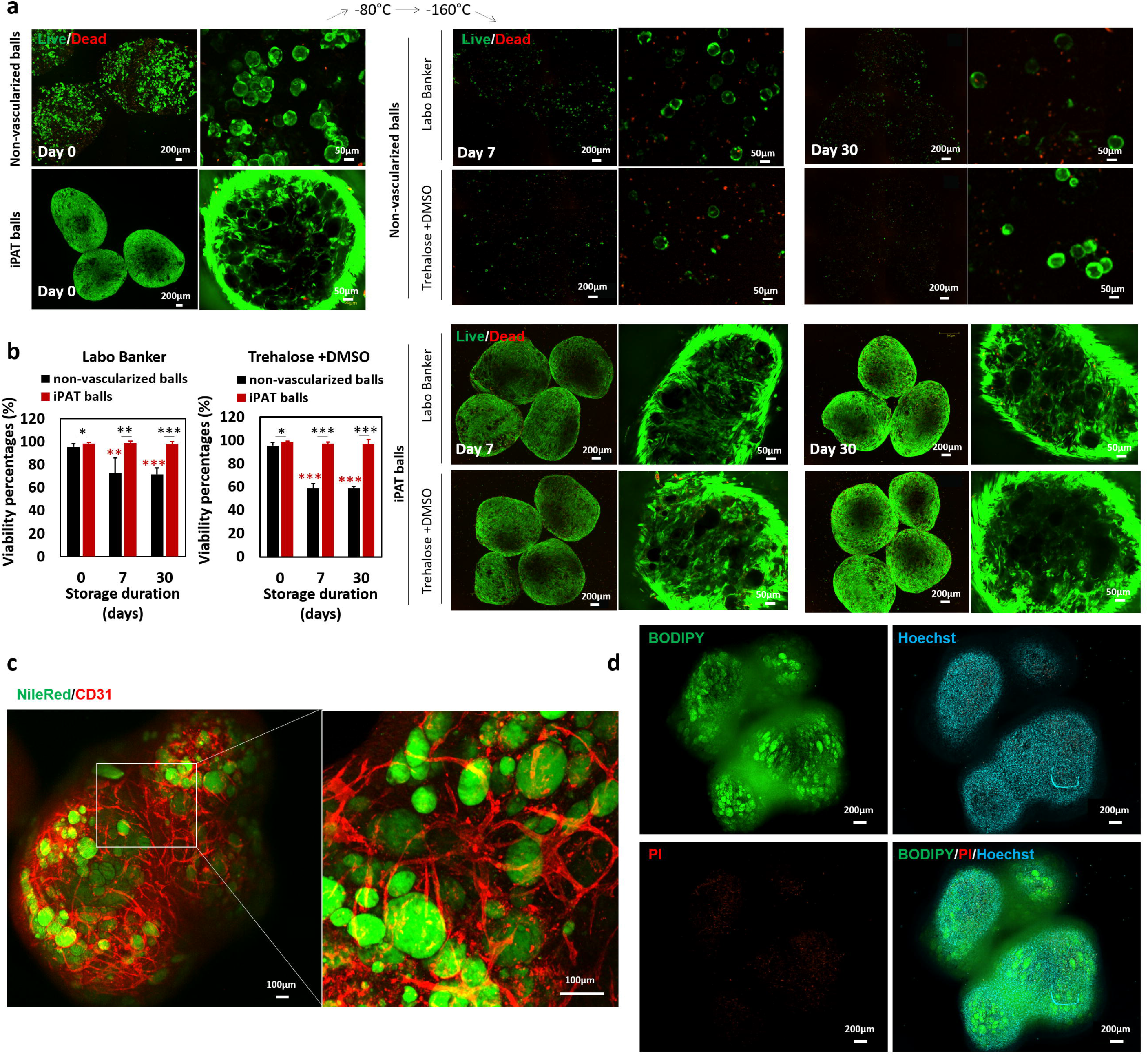
Cryopreservation of the iPATs tissues and the non-vascularized adipose tissues. After 7 days of culture, both ball tissue types were frozen in cryopreservation media. (a) Representative images of Live/Dead assays on the iPATs and non-vascularized adipose ball tissues cryopreserved in either LaboBanker or Trehalose/DMSO/FBS cryopreservation media. The viability was monitored at day 0, and after 7 and 30 days of freezing storage. (b) Quantitation of the viability percentages through the storage time, cryopreservation media and conditions. All graph results show means□±□s.d. Measures were performed on n=4-6 images per samples. \**p*<0.05, \*\**p*<0.01 and \*\*\**p*<0.001. Red asterisks are compared with the non-vascularized adipose balls at day 0. (c) Representative image of NileRed and CD31 immunostaining of several merged iPAT balls stored for 30 days and cultured 7 days after thawing in the same well. (d) Representative images of functional assessment by BODIPY™ 500/510 C1, C12 equivalent of fatty acids uptake in mature adipocytes after insulin induction, counterstained by Hoechst and Propidium Iodide (PI). n=4 balls/well and 3 samples per condition were performed.

## Discussion

We have demonstrated a novel tissue-engineering approach for improving the human adipose tissue graft survival after implantation, using iPAT tissues. Our method of mixing mature adipocytes, ADSC and HUVEC with CMF in a fibrin gel allowed the reconstruction of these vascularized adipose ball tissues which can be injected in a non-invasive way. This enables a customized final implanted volume depending on the number of ball tissues injected which is a significant advantage for clinical use. Unlike classic liposuctioned adipose tissue reinjection or engineered adipose tissues without prevascularized structures, vascularized adipose ball tissues demonstrated a higher host’s vasculature connection for both blood vessels and lymphatic networks. This may come from the added ADSC and HUVEC to the mature adipocyte tissues, because these cells can improve the revascularization through excreted angiogenic factors, provided directly by ADSC or by the blood supply formed by the HUVEC, in addition to the environmental support ^56^. The host contact allows blood flow to provide nutrients, oxygen, growth factors, hormones or cytokines important for the adipose tissue homeostasis, where the growth of blood vessels generally precedes the adipocyte differentiation and maturation ^57,58^, or is remodelled depending on the adipose tissue shrinkage or enlargement to ensure the proper energy balance of the body ^57,59^. Engineered prevascularized tissues are thus the key to accelerating the connection with the *in situ* vessels and supporting their survival and functionality ^60^. The lymphatic system is responsible for the transport of fluid, cells and lipid, helping to coordinate interstitial fluid, lipid homeostasis and immune responses. Lymphatic vasculature has just recently been recognized as a critical modulator of metabolism and obesity in particular, with a strong clinical correlation between lymphatic dysfunction and adipogenesis, leading to adipose tissue inflammation by the recruitment of macrophages and fibrosis formation ^61,62^.^61,62^. While further studies are needed to better understand the impact of the lymphatic vessels in our study, these results may provide leads for future improvements of adipose graft survival.

The adipose tissue innervation and the brain–adipose neural communications have also been investigated only recently, providing new insights into innervation patterns, neuro-immune interactions, and regulation of nerve plasticity in adipose depots. The way a nerve supply can regulate the whole-body energy balance and its metabolic health may also be of importance for improving the fat graft maintenance. For example, the release of the neurotransmitter norepinephrine is critical for adipose metabolic processes such as lipolysis, adipogenesis, and browning ^51^. Regeneration of the neural vasculature can also improve the sensory recovery of breast reconstruction patients, improving their quality of life, which is one of the current advantages of the DIEP flap method with neurotization ^23^, despite being difficult to perform.

Concerning the already published engineered adipose tissues methods for soft tissue regeneration, studies have generally used different types of biomaterials (synthetic or natural) mixed or not mixed with cells, and then checked the adipogenesis and angiogenesis found inside the tissues after implantation, without necessarily reconstructing a blood vessel vascular structure *in vitro* before the implantation (summary of published results in Table 2). The vast majority of these studies also do not verify the graft volume or weight maintenance rate after a long-term implantation. Most of them have focused more on the adipogenesis or the neovascularization achieved *in vivo*. From these results, it can be noted that the methods that stand out are those including natural scaffolds, reproducing in a more physiological way the adipose tissue ECM. In our case, the method for quantifying the survival volume differs from that generally used which consists of measuring the volume of the scaffold before implantation. Here, the adipose ball tissues include the weight of the culture medium which was difficult to dry, while this medium should diffuse in the mouse after injection and additionally, a merging of the balls occurs which should lead to a subsequent lower volume. If we thus compare the weight of the different types of ball tissues (Supplementary Figure 5b) and use it to calculate the survival percentage with respect to their weights, it appeared low with only 54% remaining after three months in the vascularized adipose ball tissues. Knowing that the major part of the adipose tissue is composed of mature adipocytes, in which >95% of adipose lipids are triacylglycerols, the density of triolein (0.915 g/mL) can be used to calculate the density of the human fat tissue ^63^.^63^. We can thus assume that the injected 100µL volume is about 90mg of weight. This therefore means that the weight survival percentage would be finally higher, at 73% after three months for the vascularized adipose ball tissues. The high scale bar observed at the three months’ data relies also on the patient’s variability, the three patients used for isolating the cells for the *in vivo* implantation had very different BMIs of 28.5, 36.6 and 19.5. In particular, obese patients often contain large inflamed mature adipocytes, which results in the lower harvest of fat stem cells which can be associated with inferior graft survival. The ADSC ability to differentiate can also be impaired depending on the patient’s BMI ^64^. However, the highest BMI patient here was the one associated with the best survival results. Our method seemed to allow the enrichment of ADSC and blood vessel vascularization even in patients expected to provide lower results in long-term approaches.

**Table 2:**
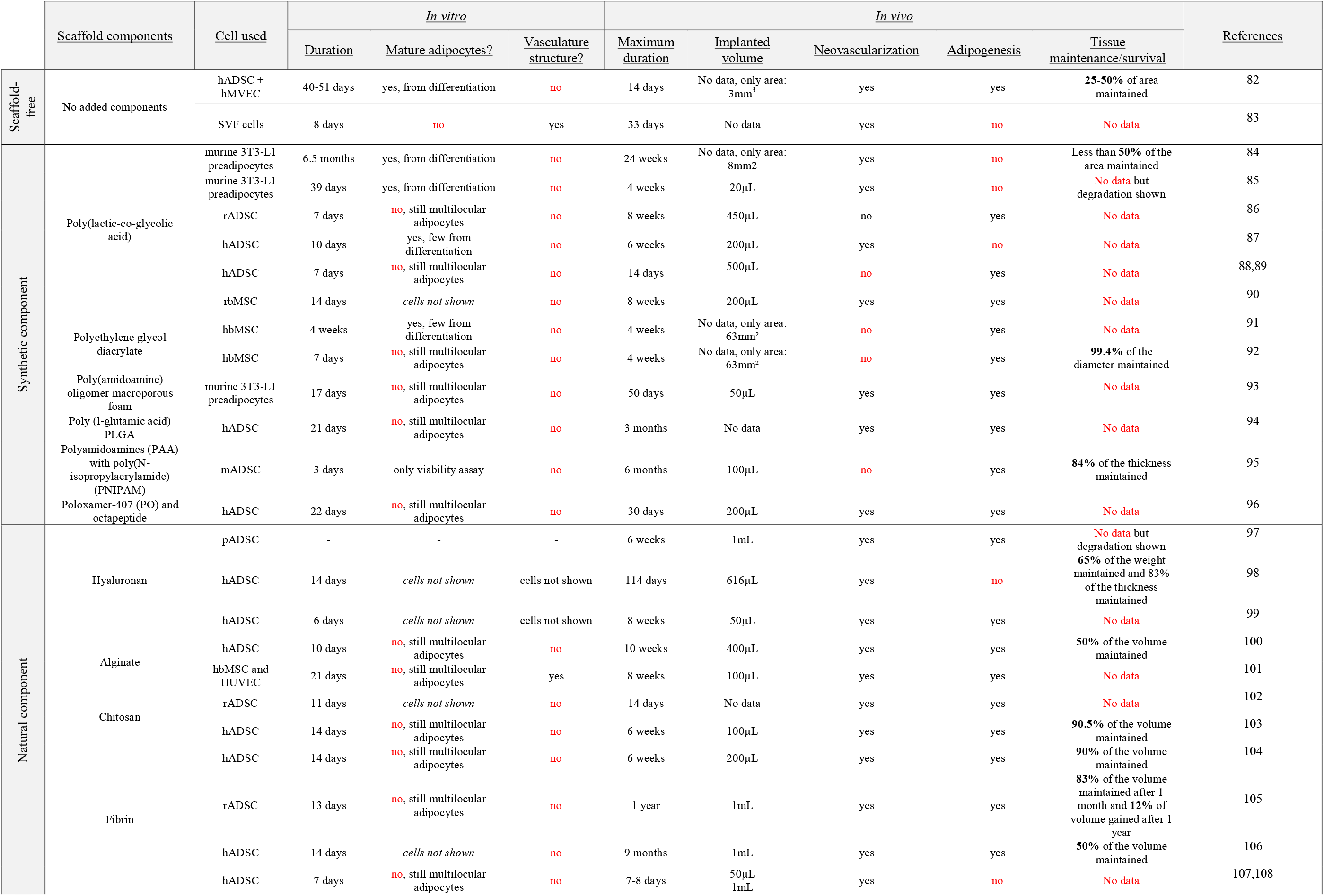
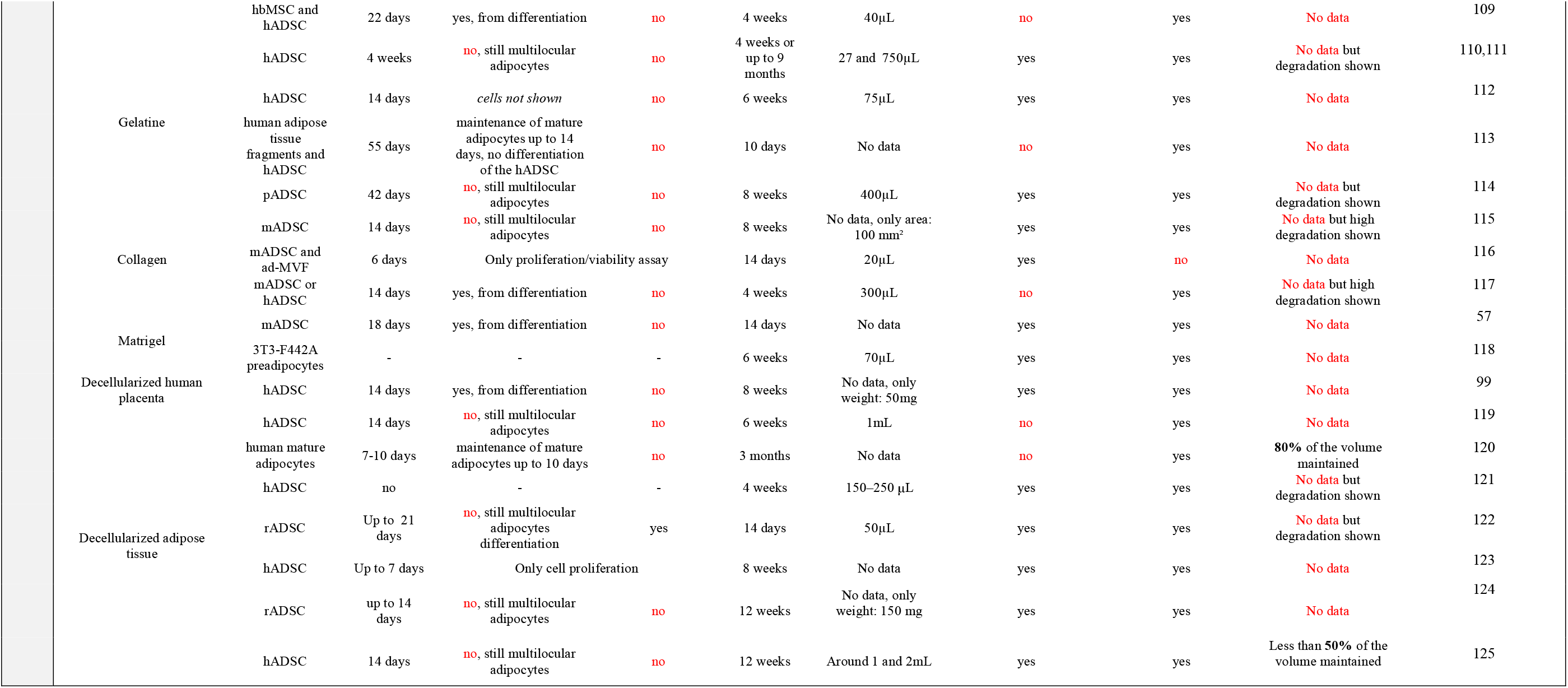
Non exhaustive summary of the recent published studies about implanted engineered adipose tissues. <u>Abbreviations:</u> SVF (stromal vascular fraction), hMVEC (human microvascular endothelial cells), hADSC (human adipose-derived stem cells), rADSC (rat adipose-derived stem cells), mADSC (mouse adipose-derived stem cells), pADSC (porcine adipose-derived stem cells), ad-MVF (adipose tissue□derived microvascular fragments), hbMSC (human bone marrow stromal cells), rbMSC (rabbit bone marrow stromal cells), HUVEC (human umbilical vein endothelial cells).

For the formation of endothelial tubular structures, these ADSC are of great importance ^65^ (as also shown in Supplementary Figure 1a), along with the ECM stroma interactions for directing the vessel growth ^38,66^, and lipoaspirates allegedly rich in ADSCs are already in clinical use ^67–69^. In implantation studies, ADSC were already found to be associated with the promotion of early revascularization leading to an 80% retention volume of the autologous fat graft ^27^. ADSC help the graft survival through their ability to stimulate not only angiogenesis^12,70,71^, but also lymphangiogenesis and by their anti-fibrogenic and anti-inflammatory effects ^72,73^. The roles of ADSC and mature adipocytes are even complementary. Differentiated ADSC into preadipocytes showed an increased transcription of the angiopoietin 1 (ANGPT1) compared to mature adipocytes ^74^ and took over the role of perivascular cells *in vivo*, supporting endothelial cell survival ^75^. Preadipocytes also showed a higher expression of the vascular endothelial growth factor (VEGF) C and D, responsible for angiogenesis and endothelial cells growth, by stimulating their proliferation and migration, and are also involved in lymphogenesis ^74^. On the other hand, Curat et al. found that mature adipocytes can induce the expression of CD31 in adipose endothelial cells, most likely through their secreted leptin, which then activate the endothelial cells’ adhesion and migration ^76^. Taken together, these findings can explain the denser vascularization, as well as the induced lymphatic network colonization, found in the vascularized adipose ball tissues when mature adipocytes and ADSC are both present.

Another concern was about the maturation of the reconstructed vasculature after its implantation, which is essential for its physiological function. Without proper prior *in vitro* maturation and stabilization of the vasculature, immature microvessels can have limited potential to anastamose to the host vasculature ^77^ leading to their regression ^78^, due to their more fragile state, which forms oedema after implantation ^79^. Accordingly, Nör et al. showed that generally, one *in vitro* capillary in five was finally perfused *in vivo* and the other capillary-like structures regressed and disappeared within 21 days ^80^. In this context, our model already showed similar features with the *in vivo* adipose tissue vasculature, in terms of the blood vessels’ distances and density, as well as for their diameters (Figure 2d). This mature phenotype may definitely help the host anastomosis and whole tissue survival after implantation.

A final concern rested on the shrinkage of the graft tissue after injection which may require subsequent injections until reaching the desired final volume. The ability to store the iPAT balls with a high cell viability and functionality after thawing will enable us to overcome this obstacle by storing sufficient numbers of iPAT balls constructed from one adipose tissue liposuction isolation on the patient, allowing for later reinjections in a non-invasive way. Previous studies have shown varied and sometimes contradictory results following the storage of adipose tissues ^81^.

In addition, further steps which will focus on the scale-up process by the assistance of bioprinters and bioreactors should finally address the significant clinical needs of effective soft tissue regenerative medicine.

## Methods

### Materials

Porcine type I collagen were kindly donated from Nippon Ham (Osaka, Japan). Fibrinogen (from bovine plasma, F8630), Thrombin (from bovine plasma, T4648), bovine serum albumin (BSA, A3294), *In Situ* Cell Death Detection Kit TMR red, Phosphate buffered saline powder (PBS, D5652), collagenase from Clostridium histolyticum (Type I, C0130) and Triton-X 100 (T8787) were purchased from Sigma-Aldrich (St Louis, MO, USA). Foetal bovine serum (35010CV) were purchased from Corning (Corning, NY, USA). Rabbit anti-neurofilament antibody (ab8135) and rabbit anti-mouse CD31 antibody (ab222783) were obtained from Abcam (Cambridge, United Kingdom). Penicillin, streptomycin, BODIPY™ 500/510 C1, C12 (4,4-Difluoro-5-Methyl-4-Bora-3a,4a-Diaza-s-Indacene-3-Dodecanoic Acid, (D3823)), goat anti-rabbit secondary antibodies Alexa Fluor® 488, goat anti-mouse secondary antibody Alexa Fluor® 647, R&D systems sheep anti-human CD31/PECAM-1 antibody (AF806) and Hoechst 33324 (H3570) were obtained from Thermo Fisher Scientific (Whaltam, MA, USA). 4%-paraformaldehyde (PFA, 16310245) and mouse anti-human CD31 antibody (M0823) came from Wako Pure Chemical Industries (Tokyo, Japan). Human umbilical vein endothelial cell (HUVEC, C25271) and endothelial growth medium (EGM-2MV, CC4147) were purchased from LONZA (Basel, Switzerland). Rabbit anti-perilipin antibody (9349S) was from Cell Signaling Technology (Danvers, MA, USA) and NileRed™ compound from Tokyo Chemical Industry (TCI, Tokyo, Japan). Dulbecco’s Modified Eagle Medium (DMEM) came from Nacalai Tesque Inc. (Kyoto, Japan). D2-40 antibody was bought from Nichirei Biosciences (Tokyo, Japan) and rabbit anti-VWF antibody (bs-4754R) from Bioss (Woburn, MA, USA). Cell-Based Propidium Iodide Solution (10011234) came from Cayman Chemicals (Ann Arbor, MI, United States).

### Collagen microfibers preparation

Based on our previous study, the collagen microfibers were made from porcine collagen type I sponge. This porcine collagen is already used in medical reconstruction applications and appeared to be well-tolerated after implantation without activating the local inflammatory or systemic immune responses ^126,127^. It was first prepared by dehydration condensation at 200°C for 24 h of crosslinking ^43^. The crosslinked collagen sponge was next mixed with ultra-pure water at a concentration of 10 mg/mL (pH=7.4, 25°C) and homogenized for 6 mins at 30,000 rpm (Violamo VH-10 S10N-10G homogenizer, diameter of 10 mm and 115 mm length, AS ONE, Osaka, Japan). Then, the solution was ultrasonicated (Ultrasonic processor VC50, 50W, 20 kHz, Sonics and Materials, Newtown, CT, USA) in an ice bath for 100 cycles (1 cycle comprised 20 sec ultrasonication and 10 sec cooling) and filtrated (40 µm filter, microsyringe 25 mm filter holder, Merck, Darmstadt, Germany), before being freeze-dried for 48 h (Freeze dryer FDU-2200, Eyela, Tokyo, Japan). The obtained CMF was kept in a desiccator at room temperature.

### Isolation of mature adipocytes and ADSC from adipose tissues

Human abdominal adipose tissues from patients were isolated at the Kyoto Prefectural University of Medicine Hospital and kept at 25-32°C during the transportation until Osaka University. The tissues were first washed in PBS containing 5% of antibiotics. Then, 8-10 g of tissue were separated into fragments to fill the 6 wells of a 6-well plate and were minced to get around 1mm^3^ in size using autoclaved scissors and tweeters, directly in 2mL of collagenase solution at 2mg/mL in DMEM 0% FBS, 5% BSA and 1% antibiotics (sterilized by filtration) per well. After one hour of incubation at 37°C and agitation at 250 rpm, DMEM medium was added and the suspension was filtrated using a sterilized 500 µm iron mesh filter, before being centrifuged 3 minutes at 80g. The suspension was then washed two times in PBS with 5% BSA and 1% antibiotics and once in complete DMEM, by 3 minutes of centrifugation at 80g between each wash. For the washing steps, after centrifugation, mature adipocytes were found in the top layer, while stromal vascular fraction containing ADSC and blood cells were in the pellet. The liquid between the top layer and the pellet was thus aspirated and discarded using a long needle and a 10mL syringe between each wash. Then, the mature adipocytes layer was moved to a new tube and cells were counted in a 10µL isolated volume, by staining the nuclei during 15 minutes with Hoechst in DMEM and using a Turker Burk hematocytometer on a fluorescent microscope. On the other hand, the pellet was resuspended in DMEM for ADSC expansion by changing the medium every day during three days and then by passaging the cells when they reach 80% of confluency.

For the liposuctioned human adipose tissue control used in this study, abdominal adipose tissue was “liquefied” manually by mincing the tissue and aspirating it using a 2mL plastic syringe. Then the “liquefied” adipose tissue was centrifuged 5 minutes at 1,500 rpm to remove the broken vasculature and the blood part, following the exact same procedure performed at the Kyoto Prefectural University of Medicine Hospital after the liposuction procedure and before the liposuctioned fat reinjection in patients.

### Cell culture in the CMF gels

To construct the fat tissues, collagen microfibers (CMF) were first weighted and washed in DMEM without FBS before being centrifuged 1 minute at 10,000 rpm to get a final concentration in the tissues of 1.2%wt (Minispin, ThermoFisher Scientific, Whaltam, MA, USA). When needed, the ADSC and the HUVEC were added after trypsin detachment (both always used at passages 1-6) and centrifugation 1 minute at 3,500 rpm (Minispin, ThermoFisher Scientific, Whaltam, MA, USA) to get a final cell concentration of 4×10^6^ ADSC/mL and 2×10^6^ HUVEC/mL. The pellet containing CMF, ADSC and HUVEC was then mixed with the fibrinogen solution (to get a final concentration at 6mg/mL, stock solution was prepared in DMEM 0%FBS 1% antibiotics, filtrated using a 0.2µm filter) and the thrombin solution (to get a final concentration of 3U/mL, stock solution was prepared in DMEM 10%FBS 1% antibiotics, filtrated using a 0.2µm filter). Finally, the mature adipocytes were added at a final concentration of 4.5×10^6^ cells /mL and the tissues were directly seeded in 24 well plate transwells (60µL gel volume, 0.4µm polyester membrane, Costar 3470, Corning, Corning, NY, USA) or in 96 well ultra-low attachment plate (5 to 15µL volume, EZ-BindShut™II round bottom, Iwaki, Yoshida, Japan) (see Figure 1). The gelation occurred during 15 minutes in the incubator at 37°C, then the transwells were moved in a 6 well plate using self-made adaptors and with 10mL of medium, or 300µL of medium were added in the 96 well ultra-low attachment plate and the ball tissues were moved in a 24 well plate (EZ-BindShut™II flat bottom, Iwaki, Yoshida, Japan) with 2mL of medium using a spatula (one ball tissue per well to avoid the merging of the balls together). DMEM was used for the adipose tissues or EGM-2 medium + 10µg/mL of final concentration of insulin for the vascularized adipose tissues. The culture medium was renewed every 2-3 days up to 7 days of culture.

For the preliminary experiments, the determination of CMF toxicity was performed using 5%wt CMF only, without fibrin gel, mixed with 10×10^6^/mL human mature adipocytes. The cells components comparison was done using the general CMF concentration as seen above. The scaffold components comparison was assessed by removing the fibrinogen and thrombin or the CMF components, or by increasing the CMF concentration. The cell ratios comparison compared the results using twice more mature adipocytes (8×10^6^ cells/mL) and adding or not insulin at 10mg/mL final concentration in the EGM-2 medium.

### Balls tissues implantation in mice

For each experiment, 6-7 immunodepressed mice (NOD/Shi-scid,IL-2RγKO Jic, NOD.Cg-Prkdc^scid^ Il2rg^tm1Sug^/ShiJic, In-vivo Science Inc., Kawasaki, Japan) had 2 samples implanted on their skin backs. On each side, 111 balls (=100µL final volume, knowing that each ball has a volume of around 0.9 µL, the calculation was made using the mean diameter) were manually put using a spatula in an Eppendorf tube and mixed with 100 µL of 20 mg/mL fibrinogen solution in DMEM, containing 10 U/mL of thrombin final concentration. This fibrinogen concentration was supposed to be rapidly degraded after implantation while allowing enough time for the balls to merge^128^ to form the final size tissue. After 15 mins gelation, the obtained gelated tissues containing the balls were implanted in an injected-way beneath the dorsal skin of the mouse through minimal incision with a small spatula. The incision was then closed with one 6-0 nylon suture. For the liposuctioned fat tissues, 100 µL was directly injected below the mice skin using an 18G needle. 3-4 mice containing all the different combinations of the 2 samples were sacrificed after 1 month to isolate the samples, and the same procedure was done after 3 months for the remaining 3-4 mice. Following extraction, the tissues were detached from the skin and their weights were measured before being fixed in 4% PFA and being embedded in paraffin block for immunohistochemistry evaluations.

### Histology and Immunohistochemistry

Samples were fixed in 4% PFA overnight at 4°C, followed by standard dehydration before embedding in paraffin wax. Sections (5 µm) were cut using a microtome (Thermo Fisher Scientific, Whaltam, MA, USA) and mounted for hematoxylin and eosin (H&E) staining, Elastica van Gieson (EVG) staining, or immunohistochemistry assessments. For the immunohistochemistry, briefly, after deparaffinization and rehydration, the sections were first antigen-activated using a citrate buffer (0.01M, pH 6.0) at high temperature treatment during 10 minutes at 121°C. Then, the sections were blocked during 20 minutes at room temperature in 1% BSA in PBS to minimize non-specific staining. First antibody (anti-Perilipin, CD31, vWF, neurofilament or D2-40) were added in BSA 1% and incubated overnight at 4°C. Samples were finally incubated with the secondary antibodies Alexa Fluor 488 or 647 (anti-mouse, anti-rabbit or anti-sheep depending on the first antibody), at room temperature in the dark for 30 minutes. Nuclei were counterstained with Hoechst for additional 10 minutes before the slides mounting. The images were captured using an FL EVOS Auto microscope (Thermo Fisher Scientific, Whaltam, MA, USA) for the colour samples and observed using an epifluorescence microscope (Confocal Quantitative Image Cytometer CQ1, Yokogawa, Tokyo, Japan) for the fluorescent samples keeping same exposition time and excitation power for each section. The analysis of the areas and fluorescence intensities were performed using ImageJ software (Fiji, version 1.52s, National Institute of Health, Bethesda, MD, USA). For the quantitation of the immunostained area, it was normalized by the full area of the tissue. Concerning the perilipin staining, the total fluorescence intensities in the centre of the graft, from 200µm to the edges of the tissues, was quantified. For each sample, 3-6 tissues sections were measured and averaged, 2-3 tissues per condition and per time, for the 3 independent experiments.

### Viability assessment

The viability of cells was quantified using the Live/Dead® viability assay kit (Molecular Probes®, Thermo Fisher Scientific, Whaltam, MA, USA). After one PBS wash, the tissues were stained with calcein and ethidium homodimer-1 during 45 mins at 37°C in the dark and then imaged using epifluorescence Confocal Quantitative Image Cytometer CQ1. Z-stack with the same steps and using maximum intensity projection was performed keeping same exposition time and excitation power for each sample. ImageJ software was used for the analysis of the projections, calculating the percentage of living cells.

### Immunofluorescence imaging

Tissues were fixed with 4% paraformaldehyde solution in PBS overnight at 4°C. Samples were permeabilized in 0.05% Triton X-100 in PBS for 7 minutes and incubated one hour at room temperature in 1% BSA in PBS to minimize non-specific staining. First mouse anti-human CD31 antibodies were added in BSA 1% and incubated overnight at 4°C. Finally, samples were incubated with the secondary anti-mouse antibodies Alexa Fluor 647, at room temperature in the dark for 2 hours. Nuclei were counterstained with Hoechst. The samples were rinsed in PBS and observed using epifluorescence microscopes (Confocal Quantitative Image Cytometer CQ1 and Confocal Fluoview FV3000, Olympus, Tokyo, Japan). Intracellular lipid accumulation was stained during 30 minutes using NileRed™ compound diluted in PBS, with nuclei counterstained by Hoechst.

### Lipids vesicles and lumens diameters observation and measurements

The diameters of the adipocytes lipids vesicles were measured on the perilipin immunostained images of the tissues, and the distance between the blood vessels as well as their diameters on the CD31 immunostained images, using ImageJ software. The image from inside one lumen was made using surface reconstruction on 3D volume images with the Imaris software (ver. 9.2.1, Version 9.2.1, Bitplane, Belfast, UK).

### Cell death assessment on histology slides

The cell death induced by either apoptosis or necrosis was assessed by the *In Situ* Cell Death Detection Kit TMR red, a TUNEL quantitation kit which was used to quantify the DNA alteration-induced cell death from both apoptosis and necrosis pathways^129^. After deparaffinization and rehydration, the sections were first treated by citrate buffer (0.01M, pH 6.0) at high temperature treatment during 10 minutes at 121°C, following the provider protocol. Then, a labelling solution of the DNA strand breaks by terminal deoxynucleotidyl transferase (TdT) which catalyses the polymerization of labelled nucleotides to free 3’-OH DNA ends (TUNEL-reaction) was applied for 1 hour at 37°C before being washed and the tissues sections were mounted. The TMR red labelled nucleotides, incorporated in nucleotide polymers, were detected keeping same exposition time and excitation power for each section, and quantified using epifluorescence microscopes (Confocal Quantitative Image Cytometer CQ1 and Confocal Fluoview FV3000) and the ImageJ software. The total fluorescence intensity of the TMR red was normalized by the total nuclei fluorescence intensity per sample for the same area. The result was then expressed in percentages of the positive sample.

### Balls tissues cryopreservation assay

For the assessment of the cryopreservation of the balls tissues, non-vascularized adipose balls and iPAT balls were seeded and cultured for 7 days in the same way than for implantation purpose. On these two types of tissues, the viability was monitored (day 0) using a Live/Dead assay during 45 minutes incubation as explained above on 3 different balls per condition. 4 balls of each condition were then added in cryopreservation media at 4°C: the ready-to-use LaboBanker (TOSC, Tokyo, Japan) or a mixture of 6% Trehalose + 4%DMSO + 90%FBS, already published as suitable for adipose tissue freezing^130,131^. The tubes were then moved to − 80°C in a freezing container allowing controlled slow freezing (1–2°C/min, BiCell, Nihon Freezer, Osaka, Japan) for 48 hours, before being stored at −160°C in a deep freezer (Nihon Freezer, Osaka, Japan). After 7 and 30 days of storage, one cryotube of the Labobanker and one of the Trehalose mixture were thawed for each ball tissues type and the balls were washed in warm PBS before being added in 37°C DMEM during 60 minutes and then incubated 45 minutes in warm PBS containing the Live/Dead assays. The Live/Dead images were taken using the epifluorescence Confocal Quantitative Image Cytometer CQ1. For the functionality assessments, after 30 days of storage in the deep freezer, the iPAT balls were thawed and 4 balls were put in the same well of a 24 well plate (EZ-BindShut™II flat bottom, Iwaki, Yoshida, Japan) with EGM-2 + 10µg/mL of insulin and medium changes every 2-3 days. After 7 days of culture, the balls had merged together and the blood vasculature connection was assessed by CD31 immunostaining, counterstained by NileRed lipids staining and Hoechst. For the fatty acid uptake monitoring by the adipocytes, first the balls were starved during 6 hours in a DMEM medium without glucose and FBS, containing only 1% of BSA. 4□µM of the fluorescently-labeled fatty acid analog, BODIPY™ 500/510 C1, C12 was then added to the culture medium for a duration of 60□minutes with 10µg/mL of insulin to induce the fatty acids uptake, counterstained by Hoechst and Propidium Iodide (PI). Imaging was performed on the living cells, using the Confocal Quantitative Image Cytometer CQ1 at 37°C and Z-stack images with the same steps and using maximum intensity projection were performed keeping same exposition time and excitation power for each sample.

### Ethics statement

The adipose tissues were collected from Kyoto Prefectural University of Medicine Hospital (Kyoto, Japan) after abdominal adipose tissues isolation of different female donors at the ages between 42 and 71, and BMI between 19.5 and 36.6. All use was approved by the Human Ethics Committee (Approval number: ERB-C-1317-1) of the Kyoto Prefectural University of Medicine Institutional Review Board and conformed to the principles outlined in the Declaration of Helsinki.

The animals in this study were maintained at the Institute of Laboratory Animals, Graduate School of Medical Sciences, Kyoto Prefectural University of Medicine, Japan. All experiments were performed with 8-weeks-old mice. Mice were housed in a barrier facility with high-efficiency particulate air-filtered racks. The experimental protocol was approved by both Osaka University and Kyoto Prefectural University of Medicine’s Animal Research Committees (Osaka University Permit Number: 29-3-0; Kyoto Prefectural University Permit Number: M2019-153). The number of animals that was used in this study was kept to a minimum and all efforts was made to reduce animal suffering in accordance with the protocols established by the Animal Research Committee of Kyoto Prefectural University of Medicine.

### Statistical analysis

Statistical analyses were performed using EzAnova software (version 0.98, University of South Carolina, Columbia, SC, USA) by Tukey multiple comparison test (double-way ANOVA). For data that compared only one factor, one-way ANOVA was performed if the number of measures was the same, otherwise T Student test was performed. Error bars represent SD. *p* values were considered significantly different at least when p<0.05. When no marks are shown on the graphs, it means that the differences are not significant.

## Acknowledgements

The authors thank Nippon Ham Foods Ltd for their kind donation of collagen. This research was supported by a Kakenhi Grant-in-Aid for Early-Career Scientists (70838523), as well as a grant from the Japanese Ministry of Education, Culture, Sports, Science and Technology(18K09488).

