## Supplementary Material for "Injectable prevascularized mature adipose tissues (iPAT) to achieve long-term survival in soft tissues regeneration"

**Supplementary Figure Captions**

**Supplementary Figure 1: Preliminary developments for the suitable cells components and scaffold components leading to vasculature structure formation**

(a) Representative immunostaining images for CD31 of blood vessels vasculature depending on the cell types added with HUVEC (AD: mature adipocytes, NHDF: normal human dermal fibroblasts) after 7 days of culture. (b) Representative immunostaining images for CD31 of blood vessels vasculature depending on the scaffold components and concentrations. All images are representative of 3 independent samples.

**Supplementary Figure 2: Assessment of the viability of the iPAT tissues.** Representative image of the Live/Dead on *in vitro* vascularized adipose tissue in transwell. Alive cells are 85%±8. Results are for 3 independent samples, 7 pictures measured/experiment.

**Supplementary Figure 3: iPAT tissues assays for culture validation inside a fibrin gel**

Assessment of the possibility to keep vascularized adipose balls (yellow arrow) embedded in a fibrin gel (black arrow) for one additional week *in vitro*. Representative image of the blood vessels spreading observation through the fibrin gel. All images are representative of 3 independent samples.

**Supplementary Figure 4: Trial injection assay of the adipose balls through human skin.**

(a) iPAT tissues and non-vascularized adipose balls tissues were cultured for 7 days and injected through the dermis and epidermis parts of the same patient from who the cells were isolated. To facilitate the injection, the skins part were stuck on a 20mg/mL fibrin gel allowing also to keep the balls tissues during the additional culture week. The white arrow shows the injection hole site, which disappear after 7 days. (b) Representative images of NileRed staining for lipids vesicles and CD31 immunostaining for blood vessels vasculature of the injected iPAT tissues and non-vascularized adipose tissue observed from the bottom, showing the vasculature connection between the skin and the balls tissues. All images are representative of 3 independent samples.

**Supplementary Figure 5: *In vivo* mice implanted tissues and survival rate results after 1 and 3 months**

(a) All isolated tissues pictures for the three conditions and the 3 independents experiments using 3 different human fat tissue donors, after 1 and 3 months implantation in mice. (b) Tissue weights of the injected 100µL volume and tissues weight measurements after 1 and 3 months, expressed in survival percentage compared to the initial weights, for the 3 independent experiments.

**Supplementary Figure 6: Cell death assessment in the implanted tissues after 1 and 3 months**

Representative image of the TUNEL cell death staining on the three conditions after 1 and 3 months. Positive control represents the cell death associated to 10 minutes of DNAse-induced DNA alterations. All images are representative of 3 independent experiments.

**Supplementary Figure 7: Blood vasculature 1 month assessment on the implanted tissues**

Representative images of combined mouse and human CD31 immunostainings of blood vessels in the tissues for the 3 conditions after 1 month of implantation. All images are representative of 3 independent experiments.

**Supplementary Figure 8: Lymphatic vascularization 1 month assessment on the implanted tissues**

Representative images of D2-40 immunostainings of mouse lymphatic vessels in the tissues for the 3 conditions after 1 month of implantation. All images are representative of 3 independent experiments.

**Supplementary Figure 9: Peripheral neural invasion assessment on the implanted tissues** **and in the *in vitro* iPAT tissues**

(a) Representative images of neurofilaments immunostainings and nuclei in the tissues for the 3 conditions after 3 months of implantation. (b) Representative images of *in vitro* iPATs tissues after 7 days of culture, showing the neurofilaments following the blood vessels vasculatures. All images are representative of 3 independent experiments.

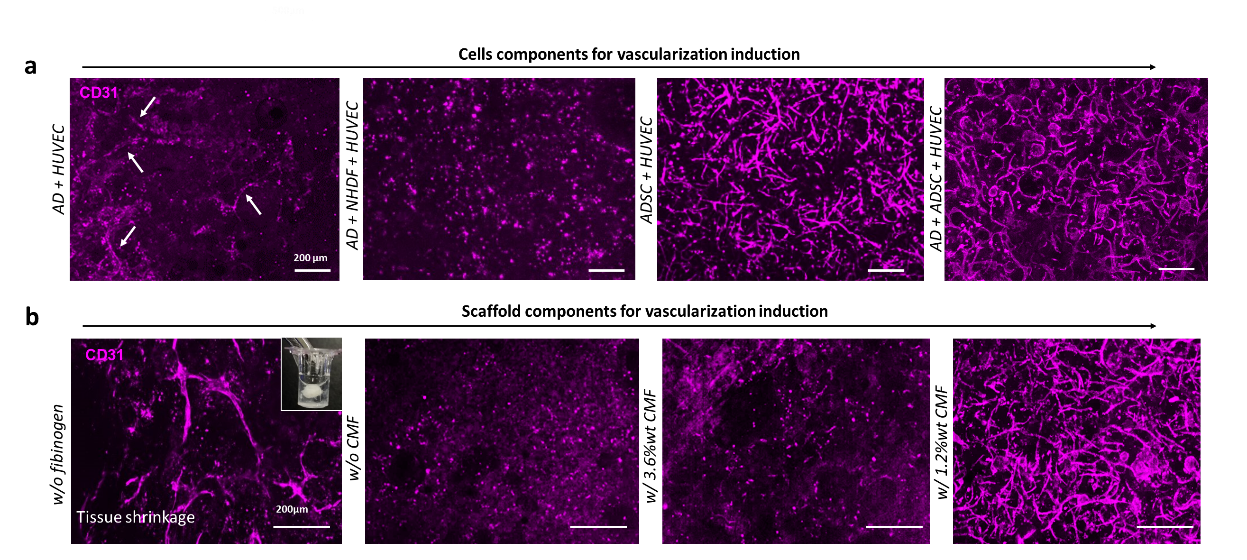

**Supplementary Figure 1**

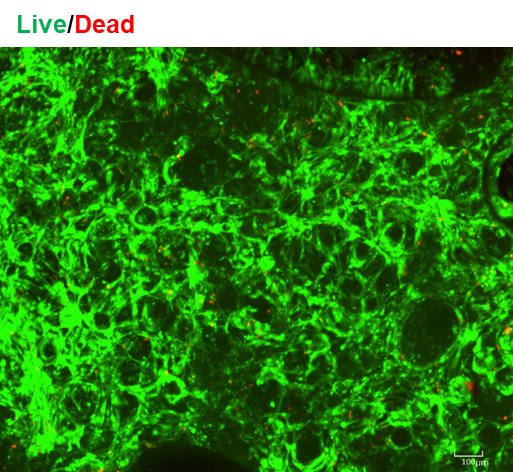

**Supplementary Figure 2**

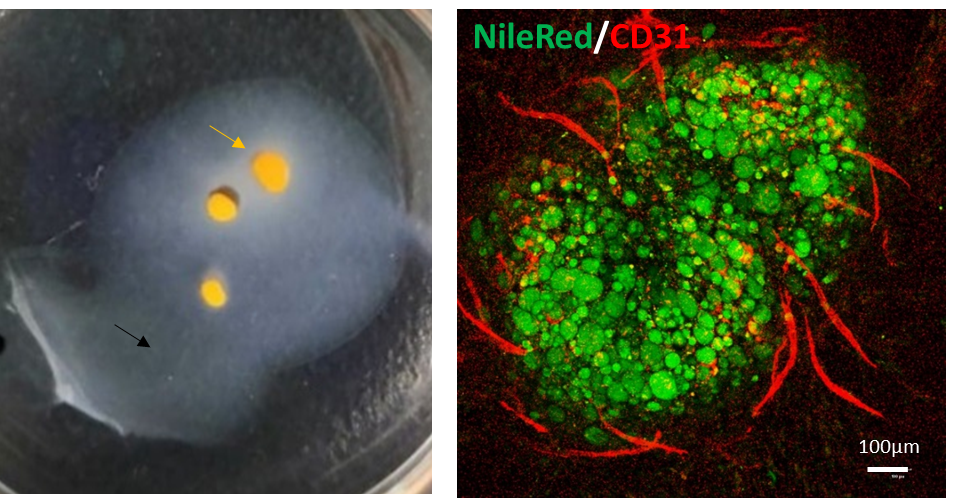

**Supplementary Figure 3**

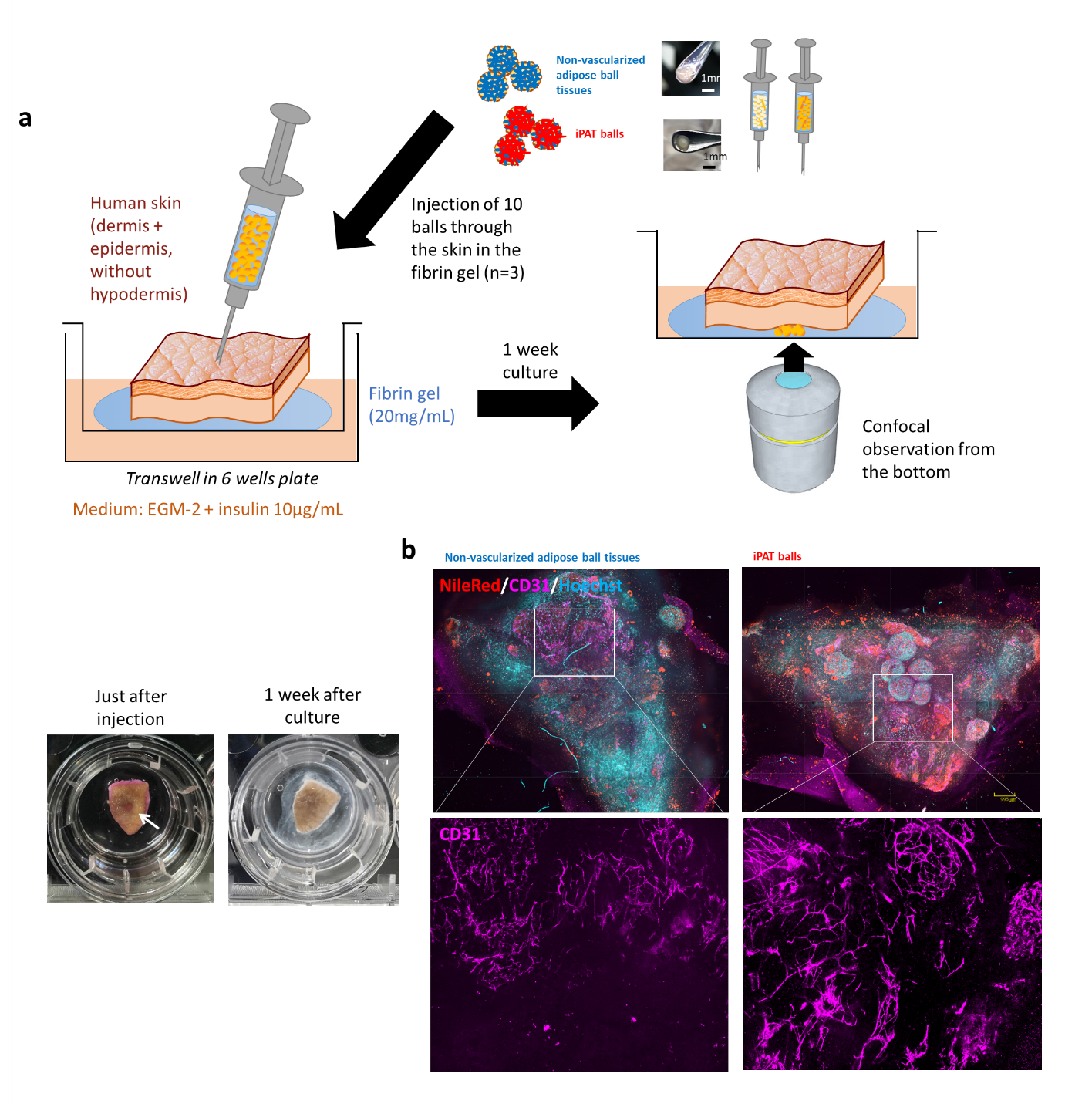

**Supplementary Figure 4**

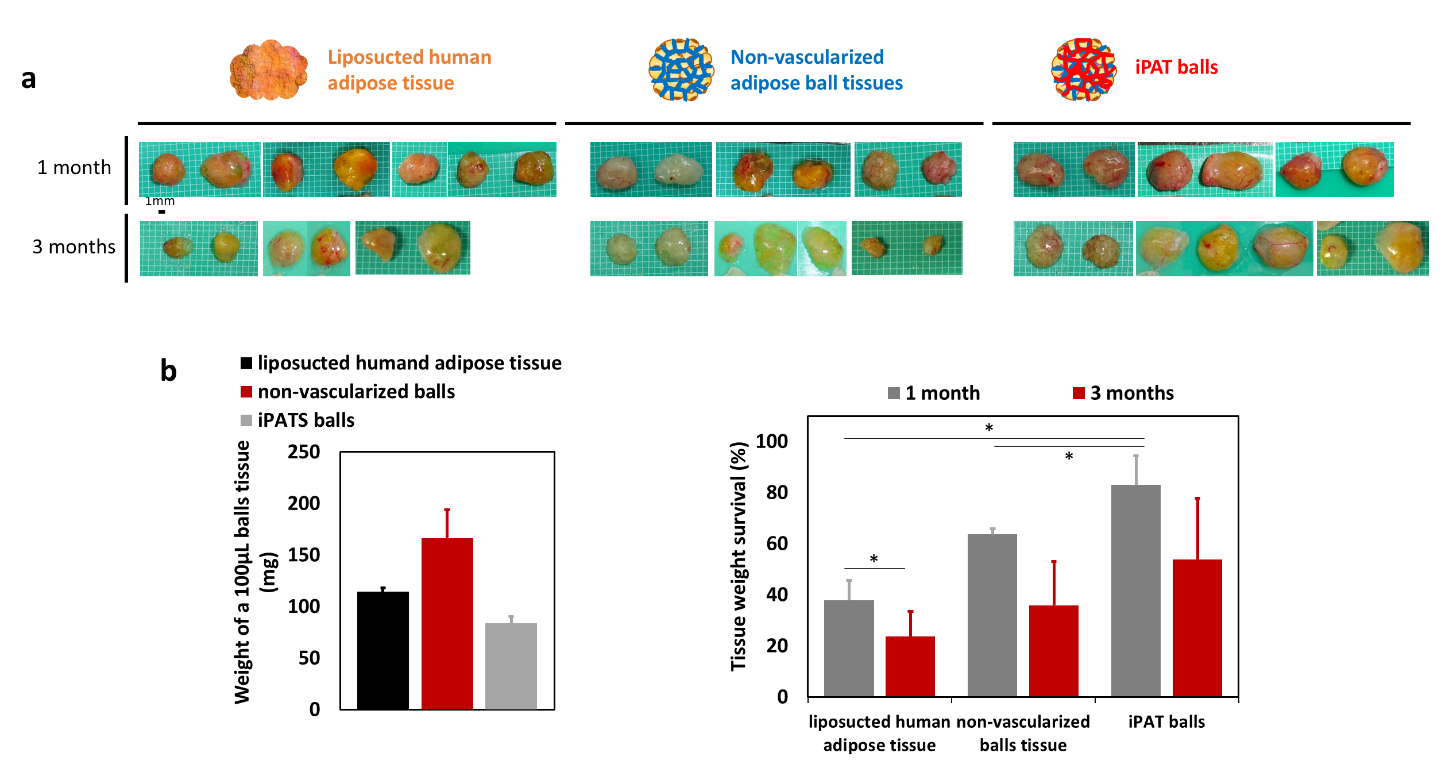

**Supplementary Figure 5**

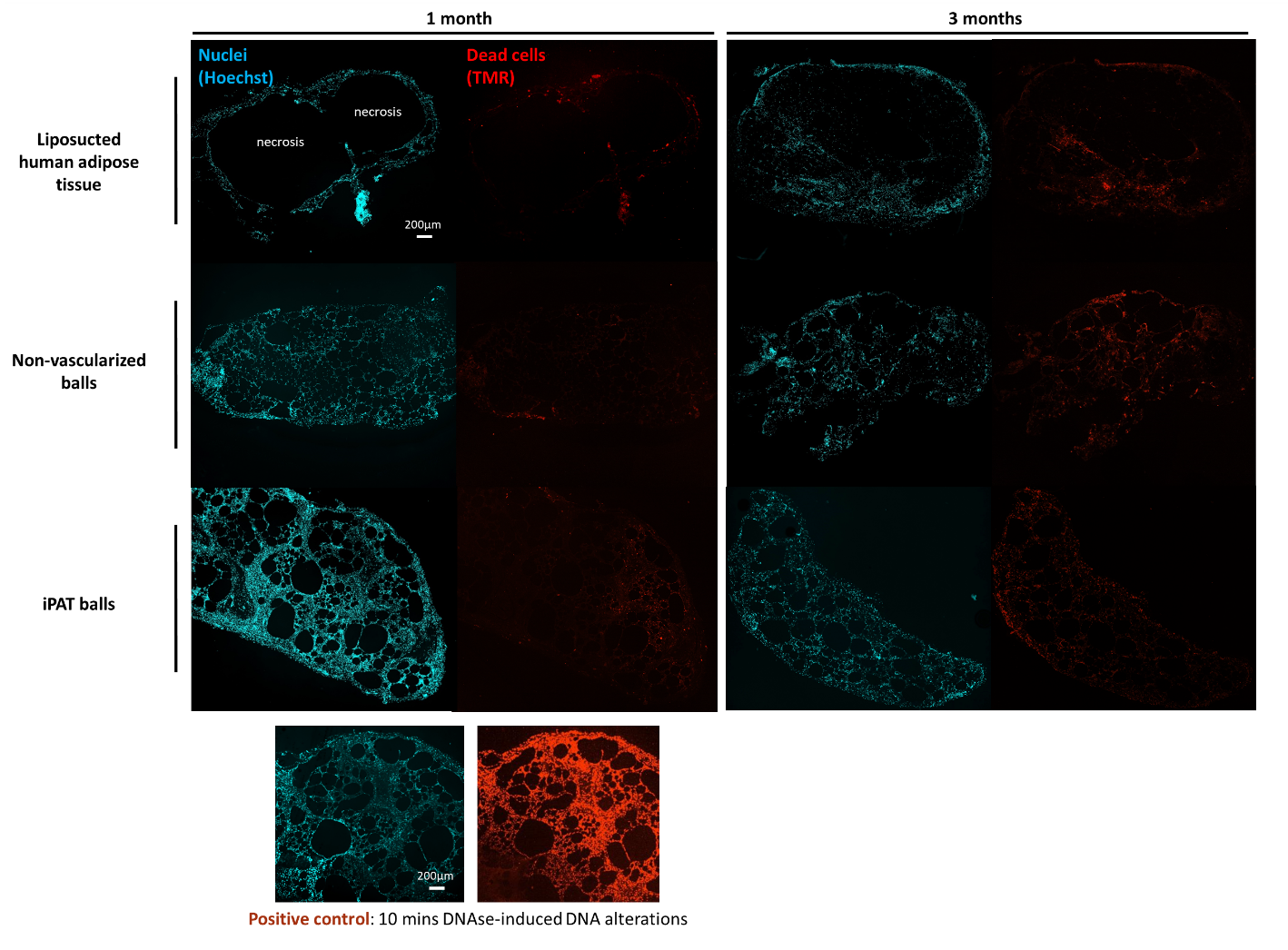

**Supplementary Figure 6**

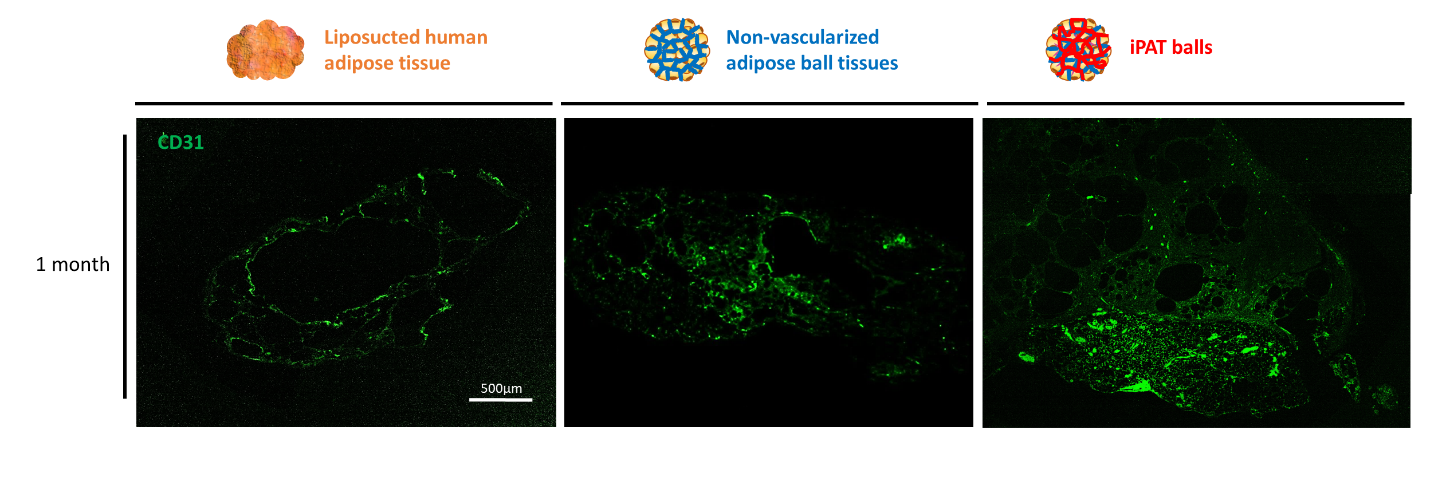

**Supplementary Figure 7**

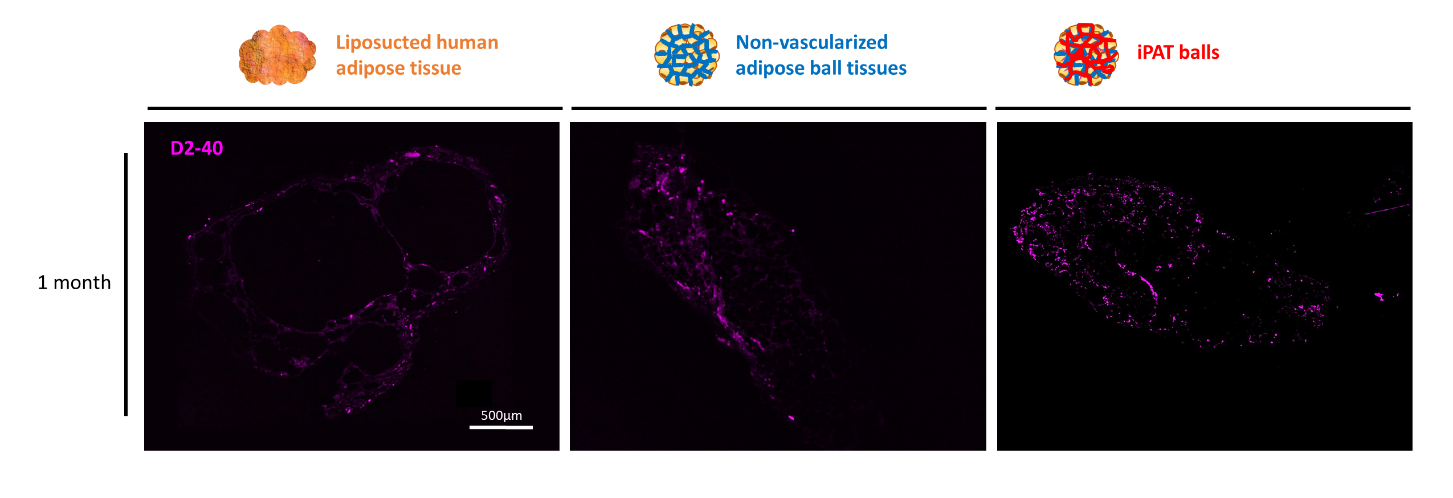

**Supplementary Figure 8**

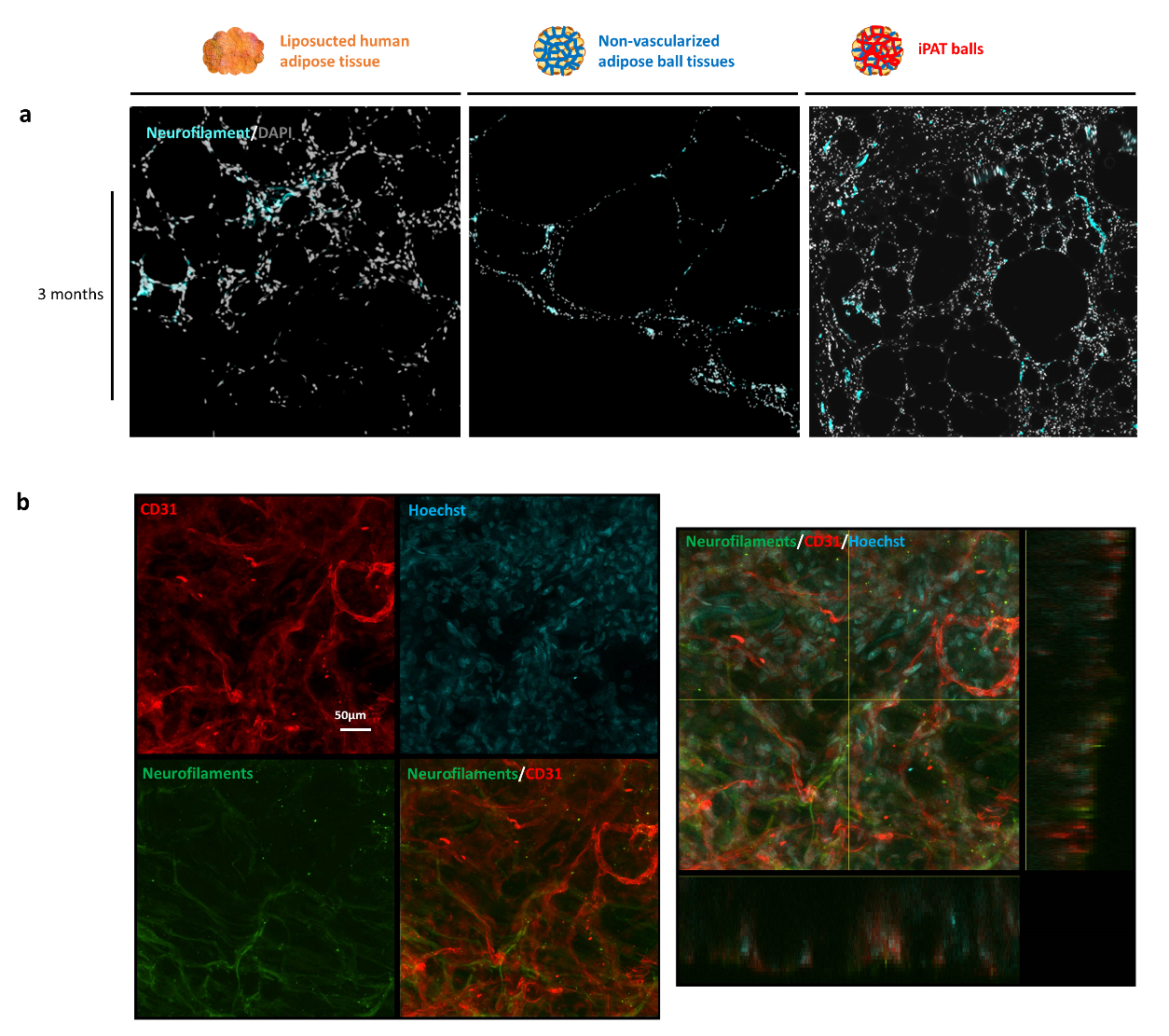

**Supplementary Figure 9**
